# Chronic exposure to low-concentration urban PM2.5 accelerates maladaptive repair after ischemic injury via mitochondrial dysfunction and lysosomal stress

**DOI:** 10.64898/2026.03.11.711056

**Authors:** Peiqi Sun, Antonio Carlos Parra, Talita Rojas Sanches, Caroline F.H. Wikuats, Loes Butter, Nike Claessen, Hans J. Baelde, Ireen Maria Schimmel, Nicole van der Wel, Georges E. Janssens, Riekelt H. Houtkooper, Frédéric M. Vaz, Joris J.T.H. Roelofs, Peter Boor, Martin Strauch, Maria de Fatima Andrade, Lucia Andrade, Sandrine Florquin, Jesper Kers, Alessia Romagnolo, Alessandra Tammaro

**Affiliations:** Department of Pathology, Amsterdam UMC Location University of Amsterdam, Amsterdam, The Netherlands; Amsterdam Cardiovascular Sciences, Diabetes and Metabolism, Amsterdam, The Netherlands; Laboratory of Basic Science in Renal Diseases (LIM-12), Division of Nephrology, School of Medicine, University of São Paulo, São Paulo, Brazil; Institute of Astronomy, Geophysics and Atmospheric Sciences (IAG), University of São Paulo, São Paulo, Brazil; Institute of Physics, University of São Paulo, São Paulo, Brazil; Department of Pathology, Systems Pathology Research group, Leiden University Medical Center, Leiden, The Netherlands; Electron Microscopy Centre Amsterdam, Amsterdam UMC, Medical Biology University of Amsterdam, Amsterdam, The Netherlands; Laboratory Genetic Metabolic Diseases, Amsterdam UMC Location University of Amsterdam, Amsterdam, The Netherlands; Amsterdam Gastroenterology Endocrinology Metabolism, Inborn errors of metabolism, Amsterdam, The Netherlands; Amsterdam UMC location University of Amsterdam, Department of Laboratory Medicine, Laboratory Genetic Metabolic Diseases, Emma Children’s Hospital, Meibergdreef 9, Amsterdam, The Netherlands; Core Facility Metabolomics, Amsterdam UMC location University of Amsterdam, Amsterdam, The Netherlands; Amsterdam Infection & Immunity, Amsterdam, The Netherlands; Institute of Pathology, RWTH University Hospital Aachen, Aachen, Germany; Department of Pathology, Erasmus Medical Center, Rotterdam, The Netherlands; Department of Internal Medicine, Nephrology division, Leiden University Medical Center, Leiden, The Netherlands

**Keywords:** PM2.5, Ischemia reperfusion injury, Senescence, Maladaptive repair, Premature aging

## Abstract

**Background:** Fine particulate matter (PM2.5), airborne particles with an aerodynamic diameter ≤2.5 μm that can penetrate deep into the lungs and enter the circulation, is increasingly recognized as a risk factor for chronic kidney disease (CKD) with long-term exposure. We previously demonstrated that high-dose PM2.5 exposure prior to ischemia–reperfusion injury (IRI) aggravates acute kidney injury (AKI). Here, we investigated how prolonged, low-concentration urban PM2.5 exposure (<15 µg/m³) affects kidney repair after AKI.

**Methods:** Six-week-old mice underwent bilateral IRI or sham surgery, followed by six months of exposure to either filtered air or ambient PM2.5 exposure in a unique exposome chamber. Kidneys were analyzed using pathomics, electron and super-resolution microscopy, immunohistochemistry, transcriptomics, and LC-MS lipidomics/metabolomics. Complementary *in vitro* hypoxia–reoxygenation and PM2.5 exposure experiments were performed in proximal tubular epithelial cells.

**Results:** Long-term PM2.5 exposure had minimal effects in sham-operated mice, including no significant changes in body weight or kidney function. Despite preserved kidney function, IRI+PM2.5 mice exhibited reduced weight gain, a marked expansion of the interstitial area, attributable to enhanced fibrosis and inflammatory responses, microvascular rarefaction, and endothelial-to-mesenchymal transition, consistent with maladaptive repair features. Proximal tubules displayed mitochondrial injury, glycolytic reprogramming, lipid accumulation, and a senescent phenotype. Energy Dispersive X-ray (EDX) microscopy confirmed PM2.5-derived elements within proximal tubules lysosomes, accompanied by lysosomal stress. Transcriptional signature–based drug screening identified nicotinamide as a compound capable of reversing PM2.5-induced metabolic alterations; *in vitro* validation confirmed restoration of mitochondrial function.

**Conclusions:** Together, these findings show that chronic post-AKI exposure to PM2.5 at levels currently considered safe by regulatory bodies drives maladaptive repair and accelerates CKD progression through mitochondrial dysfunction, lysosomal stress senescence in proximal tubules, due to local PM2.5 element accumulation.

**Translational Statement:** Acute kidney injury frequently progresses to chronic kidney disease due to maladaptive repair, yet environmental drivers of this transition remain underrecognized. Using a controlled exposome chamber, we demonstrate that chronic exposure to low, real-world concentrations of urban PM2.5 during post-ischemic recovery results in the accumulation of PM2.5-derived elements within proximal tubular lysosomes, leading to organelle dysfunction, metabolic reprogramming, lipid accumulation, and a senescence-like phenotype. Importantly, transcriptomics-based drug repurposing identified nicotinamide as a candidate compound capable of reversing metabolic dysfunction in injured proximal tubular cells subjected to hypoxia–reoxygenation and PM2.5 exposure, an effect validated *in vitro*.

## Introduction

The exposome encompasses the cumulative environmental exposures encountered throughout life that interact with genetic and physiological factors to shape organ function, disease trajectories, and aging^1^. Global analyses of more than 160,000 individuals demonstrate that environmental exposures collectively determine trajectories of healthy versus accelerated aging^2^. Among these, fine particulate matter (PM2.5, aerodynamic diameter ≤2.5 μm) is a major contributor to premature aging^3^, particularly in vulnerable populations such as individuals with prior acute kidney injury (AKI)^4–6^. Although renal function may initially recover after AKI, environmental exposures can drive maladaptive repair, leading to fibrosis and progressive decline^6^.

The proximal tubule (PT) is a primary site of susceptibility to PM2.5 exposure. *In vitro*, PM2.5 induces oxidative stress, DNA and mitochondrial damage, inflammation and activation of apoptotic signaling in proximal tubular epithelial cells (PTECs)^6–8^. Urban traffic-derived PM2.5 contains highly toxic heavy metals (e.g., lead (Pb), cadmium (Cd)), and black carbon (BC)^9, 10^. Heavy metals circulate bound to plasma proteins, such as albumin, and accumulate in kidney tissue through established PT handling pathways, including megalin- and cubilin-mediated endocytosis^11–15^. BC particles have been detected in human kidneys transplant biopsies, where their accumulation correlates with elevated urinary kidney injury molecule-1 (KIM-1)^16^. Accumulation of PM2.5 in PT may further exacerbate mitochondrial injury and metabolic stress, thereby promoting maladaptive repair. Activation of the vitamin D receptor in PT mitigates PM2.5-induced renal injury, through attenuation of oxidative stress-induced mitochondrial dysfunction, underscoring the central role of PT mitochondrial integrity in determining susceptibility^8^. Defining the mechanisms and vulnerable exposure windows underlying these effect remains an important unmet need.

We previously demonstrated that 3-months of high-dose PM2.5 exposure (600 µg/m³) before ischemia–reperfusion injury (IRI) induces mitochondrial dysfunction, inflammation and senescence in PT, leading to premature renal aging^6^. However, the impact of chronic, low-concentration PM2.5 exposure on kidney recovery after AKI remains unknown. To address this gap, we employed an exposome chamber to deliver controlled, real-world PM2.5 exposure to mice recovering from IRI. Even at concentrations largely below the WHO 24-hour guideline (15 µg/m³), six months of PM2.5 exposure after AKI induced molecular and structural alterations despite preserved renal function and the absence of overt clinical signs of CKD. PM2.5 elements accumulate within the PT lysosomal compartment, and in combination with IRI, caused PT mitochondrial dysfunction, metabolic rewiring and lipid accumulation, ultimately leading to a senescent phenotype. Transcriptomic based drug-repurposing analysis identified nicotinamide (NAM) as a candidate capable of restoring mitochondrial function *in vitro* in PTECs subjected to hypoxia/reoxygenation (H/R) and PM2.5. These findings position everyday urban PM2.5 exposure as a biologically active component of the kidney exposome and a modifiable determinant of long-term renal health.

## Methods

Detailed experimental procedures are provided in the Supplementary Methods.

### Animals and study design

Male C57BL/6J mice were randomly assigned to four groups: Sham + filtered air (FA), Sham + PM2.5, IRI + FA, and IRI + PM2.5. Bilateral renal IRI was induced by clamping the renal pedicles for 30 min. Twenty-four hours after surgery, mice were placed in whole-body exposure chambers receiving either ambient PM2.5 or FA for approximately six months. Body weight and renal function parameters were monitored longitudinally. PM2.5 samples were collected for concentration measurements, chemical characterization, and subsequent *in vitro* experiments.

### Readout measurements

At the experimental endpoint, mice were anesthetized and kidneys were collected and were harvested for downstream analyses. Renal structural and molecular alterations were assessed using a combination of deep learning–based pathomics analysis, ultrastructural and PM2.5 particle accumulation changes were evaluated by transmission electron microscopy and energy-dispersive X-ray spectroscopy. Podocyte morphology was quantified using the Podocyte Exact Morphology Measurement Procedure(PEMP). Gene expression changes were assessed by RNA sequencing followed by functional enrichment analyses. Lipidomic and metabolomic profiles were analyzed by LC–MS, and key molecular pathways were identified using multivariate and machine-learning–based approaches. Immunohistochemical analyses were performed to localize and quantify relevant cellular and molecular markers. ***In vitro* experiments** Immortalized mouse PTECs and kidney endothelial cells were used to assess cellular responses to PM2.5 exposure under H/R conditions. Senescence in pTECs was evaluated by SA-β-gal flow cytometry and p21 protein expression by Western blot. Endothelial-to-mesenchymal transition (EndMT) in kidney endothelial cells was assessed by α-SMA expression using Western blot. Mitochondrial function in PTECs was analyzed using Seahorse extracellular flux analysis. Cell viability and pharmacological modulation were evaluated using MTT assays. Where indicated, PTECs and endothelial cells were co-cultured in a Transwell system to assess paracrine interactions.

### Statistical analysis

Data distribution was assessed with the Shapiro–Wilk test. For consistency across endpoints, data are presented as mean ± SEM. Statistical testing was performed using parametric or non-parametric methods as appropriate based on data distribution. Detailed statistical procedures are described in the Supplementary Methods. A *P* < 0.05 was considered statistically significant.

## Results

### Real-world PM2.5 exposure impairs kidney repair post-AKI by promoting inflammation and fibrosis

To determine whether real-world PM2.5 exposure impairs long-term recovery after AKI, sham or IRI-challenged mice were housed for six months in FA or ambient PM2.5 in a high-traffic urban area (Fig. 1a). The mean PM2.5 concentration was 11.5 µg/m³, with 76% of the days remaining below the WHO 24-hour guideline (15 µg/m³) (Fig. 1b). PM2.5 composition consisted primarily of organic carbon (68%) and BC (12%); inorganic elements accounted for 20%, mainly consisting of sulfur (S), sodium (Na), phosphorus (P), aluminum (Al), and chlorine (Cl) (Fig. 1c).

**Figure 1.**
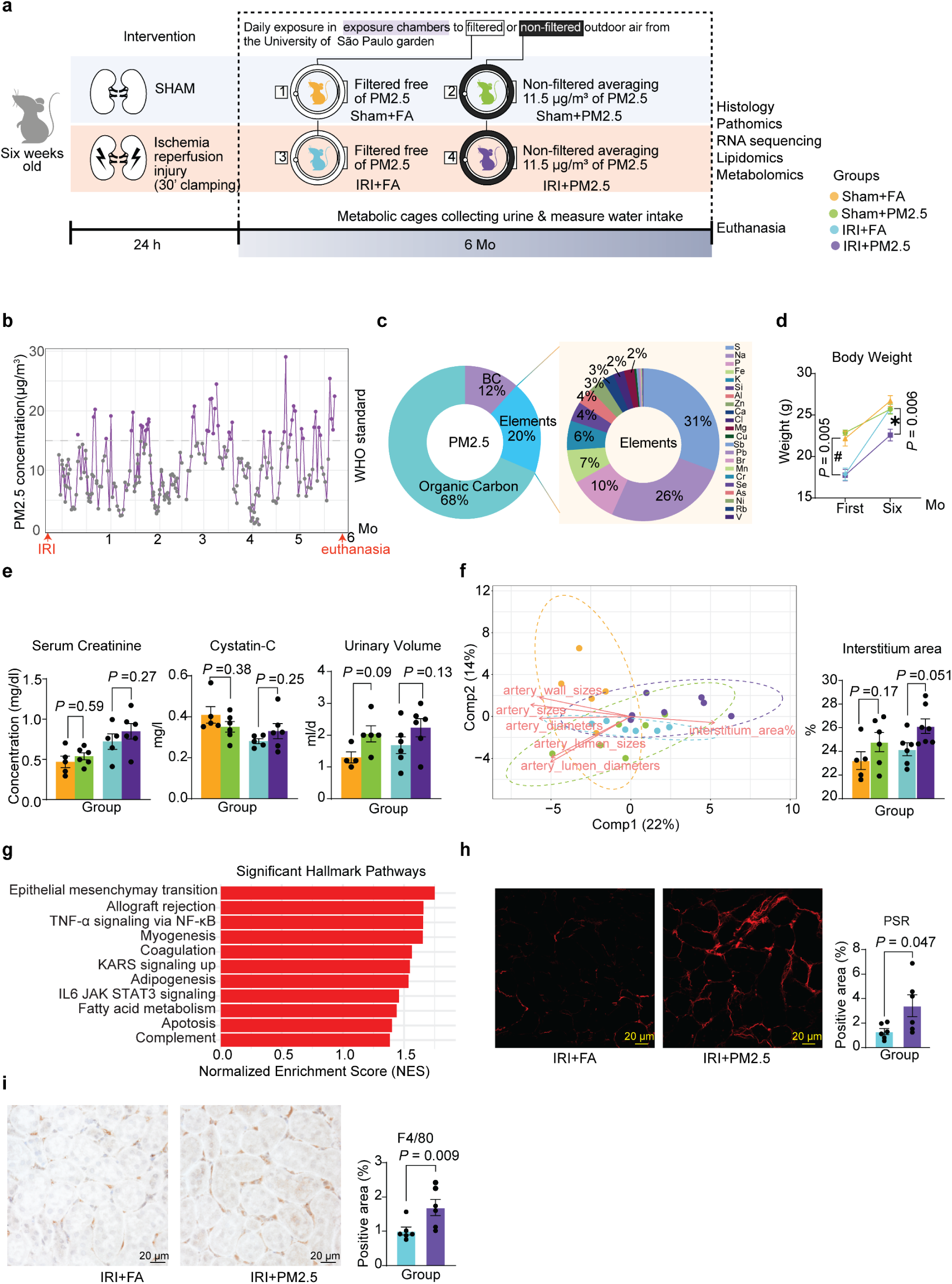
Real-world PM2.5 exposure impairs kidney repair post-AKI by promoting inflammation and fibrosis. **(a)** Schematic of the experimental design and exposure timeline. **(b)** Daily PM2.5 concentrations (µg/m³) over six-months exposure period. The purple line showed measured levels, and the dashed grey line indicated the World Health Organization (WHO) 24-h air quality limit (15 µg/m³). **(c)** Details of PM2.5 chemical composition. Left: major mass components; Right: detailed elemental composition. **(d)** Body weight at one and six months after ischemia–reperfusion injury (IRI). Data were expressed as mean ± SEM; unpaired t-test. # comparison Sham + filtered air (FA) or vs. IRI+FA at 1 month; * comparison IRI+FA vs. IRI+PM2.5 at six months. **(e)** Kidney function parameters (serum creatinine, Cystatin-C, urinary volume). Data were expressed as mean ± SEM, Serum creatinine and urinary volume were analyzed using one-way ANOVA, whereas cystatin C was analyzed using the Kruskal–Wallis test. **(f)** Pathomics analysis. Left: Partial least squares–discriminant analysis (PLS-DA) score plot showing group separation driven by morphometric features. Component 1 explains 22% of the variability, while component 2 explains 14% of the variability. Right: Pathomics scoring for interstitium area in each group (%). Data were expressed as mean ± SEM; Kruskal–Wallis test. **(g)** Hallmark pathway analysis showing significantly enriched pathways (*P* < 0.05) in IRI+PM2.5 vs. IRI+FA kidneys. Pathways were ranked by normalized enrichment score (NES). **(h)** Representative Picrosirius Red (PSR) images across groups with quantification of PSR+ area per section. Data were expressed as mean ± SEM; one-way ANOVA. **(i)** Representative image and quantification of F4/80 positive area per high-power field (HPF). Data were expressed as mean ± SEM; Kruskal–Wallis test.

IRI mice showed reduced weight gain during the first month post-injury. By six months, body weight recovered in IRI+FA mice but remained significantly lower in IRI+PM2.5 mice (Fig. 1d). Serum creatinine, cystatin-c, and urinary volume did not differ among groups (Fig. 1e). Additional metabolic cage parameters, including tubular function data such as urinary excretion of Na+ and K+, are presented in Supplementary Table S1. As functional assessments didn’t reveal marked differences, we further examined renal morphological changes using pathomics^17^. Partial least squares–discriminant analysis (PLS-DA) identified interstitium area percentage as the strongest contributors to separation of the IRI+PM2.5 group (Fig. 1f), while other features were unchanged (Supplementary Table S2).

Bulk RNA-seq revealed substantial transcriptional alterations despite limited morphological changes in sham or IRI kidneys. Chronic PM2.5 exposure induced 364 upregulated and 345 downregulated differentially expressed genes (DEGs) in sham mice and 325 upregulated and 240 downregulated DEGs in IRI mice (Supplementary Table S3). Gene Set Enrichment Analysis (GSEA) revealed positive enrichment of KRAS signaling, adipogenesis, myogenesis, and late estrogen response, together with negative enrichment of inflammatory signaling pathways in the sham group (Supplementary Fig. S1), indicating an early, subclinical response to chronic PM2.5 exposure even in health kidneys. In the IRI group, GSEA exhibited significant upregulated enrichment of epithelial–mesenchymal transition (EMT), inflammatory signaling, coagulation, apoptosis, and metabolic reprogramming pathways (Fig. 1g), while not downregulated pathways were detected. Given the stronger effects in IRI mice, subsequent analyses focused on this condition.

Expansion of the intertubular space in IRI+PM2.5 kidneys detected by pathomics can be explained by e.g. edema, inflammation, fibrosis, or a combination of these factors. Consistent with transcriptional enrichment of EMT (NES = 1.75, *P*= 0.003) and inflammation-related pathways, Picrosirius Red staining showed increased interstitial collagen deposition, and F4/80 immunostaining showed enhanced interstitial macrophage infiltration (Fig. 1h,i). Together, this exposome study show that long-term real-world exposure to PM2.5 drives coordinated transcriptomic, structural, fibrotic, and inflammatory remodeling in the post-AKI kidney, consistent with maladaptive repair and not detectable by currently used kidney function parameters.

### Chronic PM2.5 exposure exacerbates renal microvascular dysfunction, and glomerular injury in post-AKI

To identify the cellular source of the EMT-related signature, we assessed tubular EMT by vimentin immunostaining. However, no vimentin-positive tubules were detected (Fig. 2a). Because EMT gene sets can’t distinguish epithelial from endothelial cells, we next examined EndMT. Consistent with this, CD34/α-SMA co-labeling demonstrated increased spatial overlap in IRI+PM2.5 kidneys(Fig. 2b). In parallel, we observed evidence of endothelial loss. CD34 immunostaining revealed a marked reduction in both peritubular and glomerular capillary density in the IRI+PM2.5 group compared with the IRI+FA group (Fig. 2c,d).

**Figure 2.**
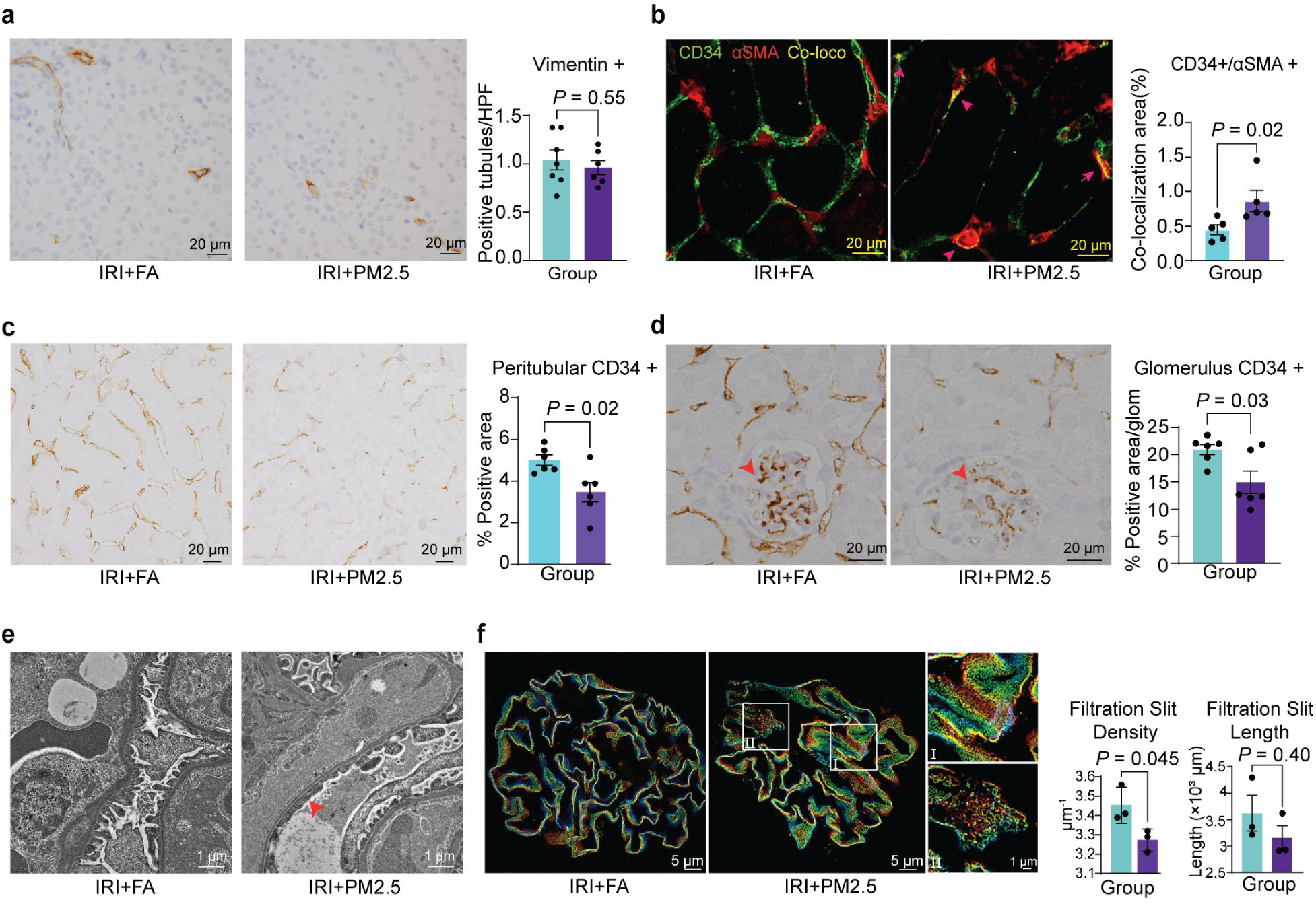
Chronic PM2.5 exposure exacerbates renal microvascular dysfunction, and glomerular injury in post-AKI. **(a)** Representative image and quantification of Vimentin+ tubules per high-power field (HPF). Barplot showed quantification of Vimentin+ tubules. Data were expressed as mean ± SEM; unpaired t-test. **(b)** Co-immunostaining of CD34+ (green) and α-SMA+ (red). Yellow regions indicated co-localization. Barplot showed quantification of co-localized area. Data were expressed as mean ± SEM; Mann-Whitney test. **(c)** Peritubular capillary density assessed by CD34 staining, excluding glomerular regions. Representative images and quantification were shown. Data were expressed as mean ± SEM; unpaired t-test. **(d)** Glomerular CD34 staining and quantification of CD34+ area from 25 glomeruli per mouse. Data were expressed as mean ± SEM; unpaired t-test. **(e)** Glomerular ultrastructure by transmission electron microscopy (TEM). Red arrows indicated areas of podocyte foot process effacement. **(f)** Podocyte slit-diaphragm morphology quantified by the Podocyte Exact Morphology Measurement Procedure (PEMP) method. (I) Representative image of a normal slit diaphragm. (Ⅱ) Representative image showing a disrupted slit diaphragm. Bar graphs showed filtration slit density (FSD) and filtration slit length (FSL). Data were expressed as mean ± SEM; FSD; unpaired t-test. FSL; Mann-Whitney test.

Because podocytes depend on the glomerular endothelium for structural support, nutrient delivery, and paracrine signaling, PM2.5-induced endothelial rarefaction may promote podocyte stress and injury. Consistently, transmission electron microscopy (TEM) revealed podocyte foot-processes effacement in IRI+PM2.5 kidneys (Fig. 2e). To quantify podocytes ultrastructural alterations, we applied the 3D-SIM super-resolution microscopy-based PEMP, which allows precise assessment of slit diaphragm integrity^18^. We observed a disrupted slit diaphragm in the IRI+PM2.5 group (Fig. 2f(Ⅱ)). Furthermore, although the total filtration slit length (FSL) remained unchanged, the filtration slit density (FSD) was reduced (Fig. 2f), supporting foot process effacement.

### PM2.5-driven microvascular dysfunction induces tubular hypoxia, mitochondrial injury, and glycolytic reprogramming

Given the critical role of peritubular capillaries in oxygen delivery to tubules^19–21^, disruption of the microvascular network may contribute to tubular hypoxia^20, 22^. Consistent with this, Gene Ontology (GO) analysis showed significantly enriched for hypoxia-related pathways among genes upregulated in the IRI+PM2.5 group (Fig. 3a). Using the iHypoxia database^23^, we identified 10 downregulated (blue) and 21 upregulated (red) hypoxia-associated genes in the IRI+PM2.5 group compared with IRI+FA (Supplementary Fig. S2a). We validated these findings by staining for vascular endothelial growth factor (VEGF), a key hypoxia-responsive protein^21^, which was significantly increased in the IRI+PM2.5 group, with stronger signals particularly visible in PT (Fig. 3b). Sustained tubular hypoxia impairs oxidative phosphorylation, resulting in mitochondrial dysfunction, and metabolic reprogramming in PTs^24^. Consistent with our previous findings that PM2.5 induces PT mitochondrial dysfunction^6^, TEM revealed exacerbated PT mitochondrial damage in the IRI+PM2.5 kidneys, characterized by swelling, cristae disruption, and the presence of electron-dense material within mitochondria (Fig. 3c,d; Supplementary Fig. S2b), suggestive of mitochondrial calcium accumulation under hypoxic conditions^25, 26^.

**Figure 3.**
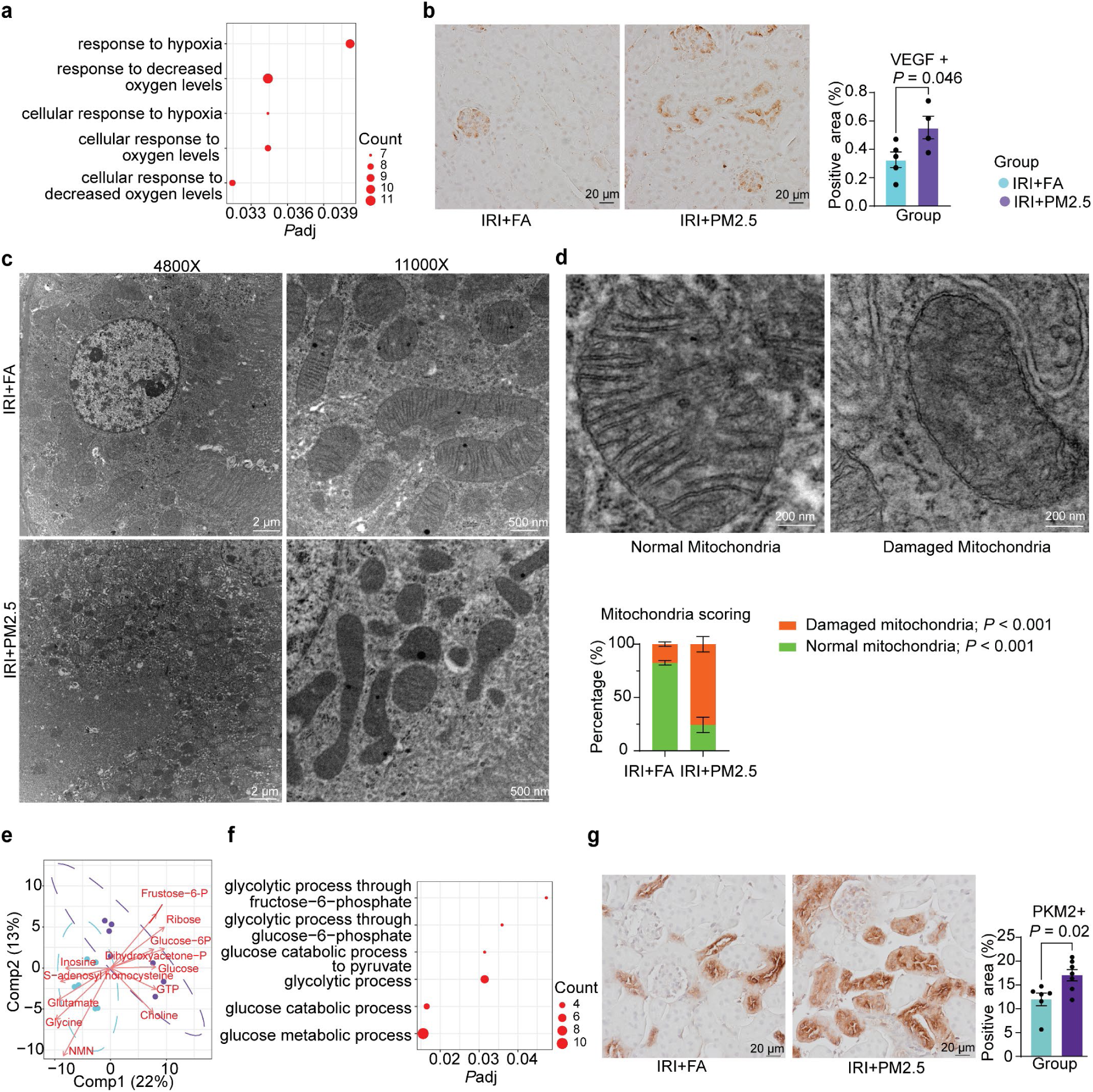
PM2.5-driven microvascular dysfunction induces tubular hypoxia, mitochondrial injury and glycolytic reprogramming. **(a)** Gene Ontology – Biological Process (GO-BP) enrichment of upregulated genes for hypoxia-related pathways. Dot size represented gene counts. **(b)** VEGF immunostaining and quantification of VEGF+ area. Data were expressed as mean ± SEM; unpaired t-test. **(c)** Representative transmission electron microscopy (TEM) images of proximal tubular mitochondria in ischemia–reperfusion injury (IRI) + filtered air (FA)and IRI+PM2.5 at 4800X and 11000X magnification. **(d)** Representative images used for mitochondrial damage scoring. Data were expressed as mean ± SEM; two-way ANOVA with Sidak’s multiple comparisons test. **(e)** Partial least squares–discriminant analysis (PLS-DA) describing the top contributing metabolites separating 2 groups. Component 1 explained 22% and Component 2 explained 13% of the variability. **(f)** GO-BP enrichment of upregulated genes for glycolysis-related pathways. Dot size represented gene counts. **(g)** PKM2 immunostaining and quantification of PKM2+ area. Data were expressed as mean ± SEM; Mann-Whitney test.

To determine whether PM2.5-driven PT mitochondria dysfunction promotes a metabolic shift away from oxidative metabolism toward glycolysis, associated with a maladaptive tubular phenotype^27^, we performed untargeted metabolomic profiling in kidney tissue. The abundance of metabolites is shown in Supplementary Fig. S2c. PLS-DA revealed partial separation between the IRI+PM2.5 and IRI+FA groups, indicating its potential alterations in metabolic profiles (Fig. 3e). The top contributing metabolites included glucose, glucose-6-phosphate (G6P), and fructose-6-phosphate. Quantitative analysis revealed a trend toward increased glucose abundance (*P*= 0.07) and a significant elevation of Glucose-6-Phosphate (*P*= 0.02) in the IRI+PM2.5 group compared with the IRI+FA group (Supplementary Fig. S2d), consistent with enhanced glycolysis. In contrast, tricarboxylic acid (TCA) cycle metabolites like citrate and succinate were unchanged (Supplementary Fig. S2c). Consistent with the metabolomics profile, RNAseq demonstrated significant enrichment of glycolysis-related pathways (Fig. 3f). However, as bulk data cannot identify the specific cell types involved, we stained for pyruvate kinase M2 (PKM2), a key glycolytic enzyme^28^, which was increased in PTs of the IRI+PM2.5 group (Fig. 3g).

### PM2.5 elements reach the kidney and accumulate in PT lysosomes, inducing lipid remodeling and lysosomal stress

Mitochondrial dysfunction may induce lipid accumulation and lipotoxic stress in PT, through fatty acid oxidation (FAO) impairment, thereby inducing failed-repair mechanisms and tubular senescence^29^. Consistent with this, TEM revealed atypical lipid accumulation in between swollen endoplasmic reticulum (ER) within the PT of IRI+PM2.5 kidneys (Fig. 4a, red arrowhead). To assess whether FAO was impaired, we first examined transcriptional signatures. GSEA didn’t show overt suppression of FAO pathways (Supplementary Fig. S3a). Similarly, Cpt1a, the rate-limiting enzyme for mitochondrial FAO, showed similar protein expression between groups (Supplementary Fig. S3b). However, GSEA of lipid catabolic processes showed significantly enrichment in IRI+PM2.5 kidneys (Fig. 4b), suggesting altered lipid degradation a possible consequence of lipid accumulation^30^. To further characterize lipid alterations, we performed semi-targeted lipidomics, identifying a total of 2,281 lipid species across 63 classes. Differential analysis revealed a distinct PM2.5-associated lipid signature, with 185 differential lipid species identified, including 155 upregulated and 30 downregulated species (*P*< 0.05; Supplementary Fig. S3c). A heatmap using a more stringent threshold (*P*< 0.01) highlighted coordinated increases in multiple lipid classes, including triacylglycerols (TG); glycerophospholipids such as phosphatidylcholine (PC), phosphatidylethanolamine (PE), and phosphatidylserine (PS); lysosome related phospholipids including bis(monoacylglycero)phosphate (BMP); sphingolipids (SM); and Hexosylceramide (HexCer)(Fig. 4c). PLS-DA and Variable importance in projection (VIP) scores, identified several BMP (e.g. BMP 36:4–36:6; 38:7–38:8), as well as SM (e.g SM(t38:0–t38:1) and HexCer(d44:3)) species among the top discriminative lipids (Fig. 4d,e). Random forest analysis confirmed BMP (36:5–36:6; 38:7), SM (t38:1, t34:1), and HexCer (t41:2; d44:3) as top-ranked features (Supplementary Fig. S3d), highlighting lysosomal lipids as key contributors to the PM2.5-associated lipid signature. These consistent with increased expression of *Gpnmb* (Fig. 5a), a marker of lysosomal stress, in the IRI+PM2.5 group. Moreover, GO enrichment analysis of DEGs revealed enrichment of early endosome–related terms (Supplementary Fig. S3e).

**Figure 4.**
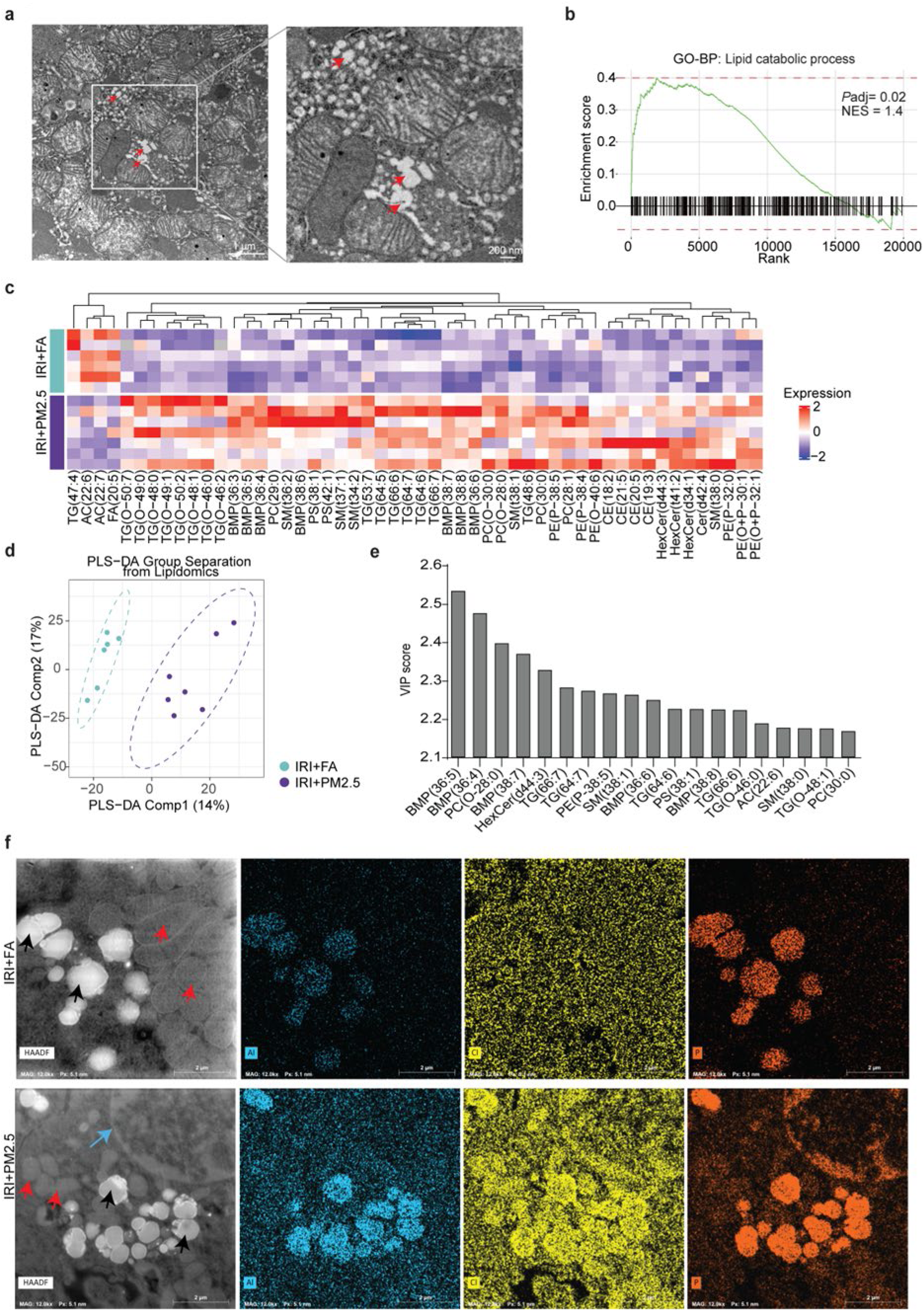
PM2.5 elements reach the kidney and accumulate in PT lysosomes, inducing lipid remodeling and lysosomal stress. **(a)** Transmission electron microscopy (TEM) images showing prominent lipid droplet accumulation in the ischemia–reperfusion injury (IRI) + PM2.5 group; red arrowheads indicate lipid droplets. **(b)** Gene Set Enrichment Analysis (GSEA) plot of lipid catabolic process from Gene Ontology – Biological Process (GO-BP) pathways. The green line represented the enrichment score, and black vertical ticks indicated the positions of pathway genes in the ranked gene list. Statistical metrics for the pathway were shown on the right. **(c)** Heatmap of lipid species with differential abundance (*P* < 0.01). Rows represented Z-score–scaled lipid molecules, and columns represented individual samples. **(d)** Partial least squares–discriminant analysis (PLS-DA) showed the lipidomics data can clearly discriminate between IRI + filtered air (FA) and IRI+PM2.5, with component 1 explaining 14% of the variability and component 2 explaining 17% of the variability. **(e)** Variable Importance in Projection (VIP) scores identifying the top lipid species contributing to group discrimination in the PLS-DA model. Lipids with higher VIP scores represented the variables driving class separation. **(f)** Energy-dispersive X-ray spectroscopy (EDX) imaging of proximal tubules (PT) lysosomes. The IRI+PM2.5 group showed clear elemental signals (Al, P, Cl) consistent with PM2.5-derived inorganic particle deposition within lysosomes. Red arrows indicated mitochondria, blue arrow indicated nuclear and black arrows indicated lysosomes.

**Figure 5.**
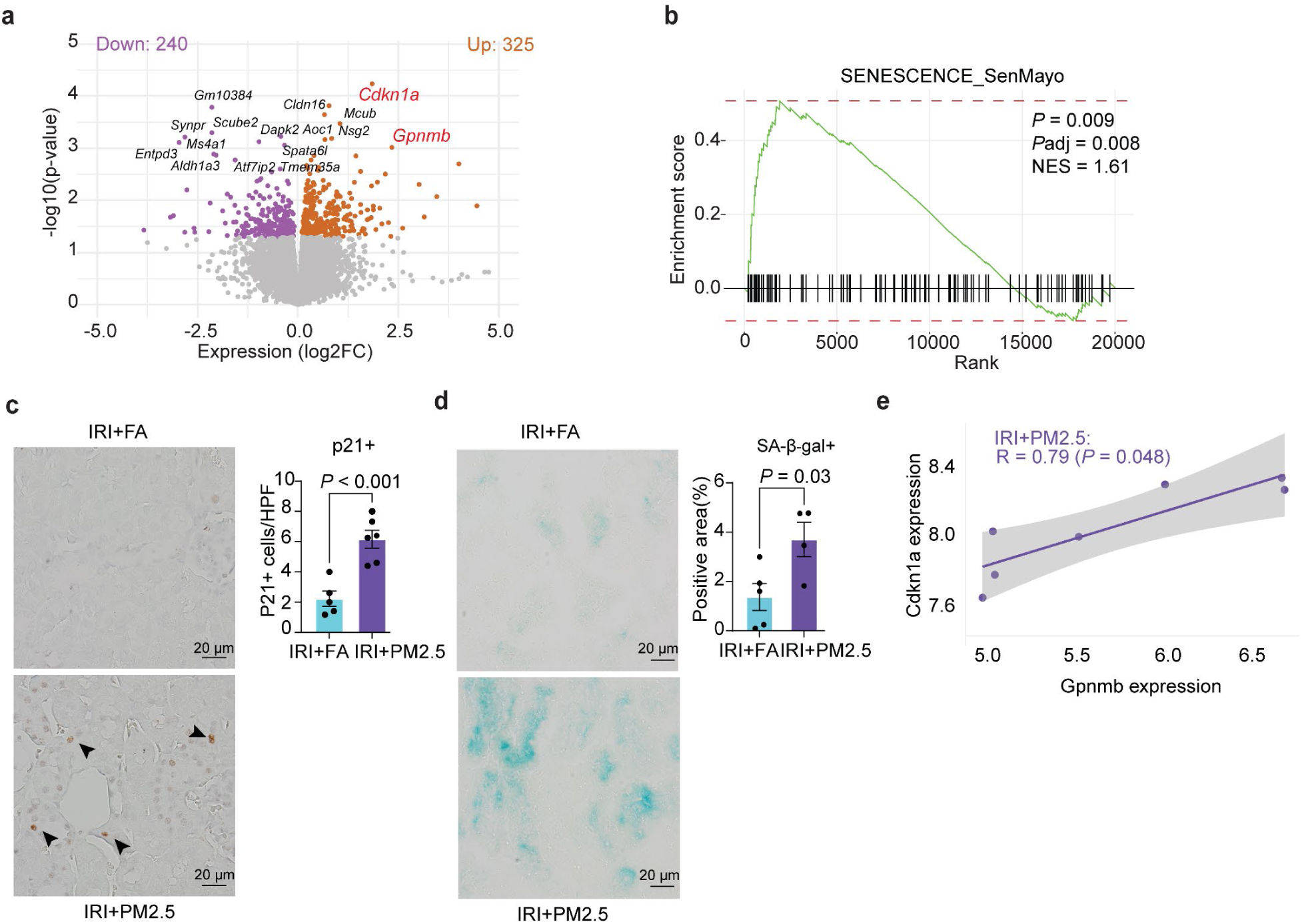
PM2.5-driven organelle stress promotes maladaptive repair through tubular senescence. **(a)** Volcano plot showing differentially expressed genes (DEGs) between ischemia–reperfusion injury (IRI) + filtered air (FA) and IRI+PM2.5 kidneys, highlighting significantly upregulated (orange) and downregulated (purple) genes. *Cdkn1a* (p21) as key senescence marker, and *Gpnmb* as lysosome stress maker, highlighted in red. **(b)** Gene Set Enrichment Analysis (GSEA) plot showing significant enrichment of the SenMayo senescence dataset. Green line indicated enrichment score; black vertical lines represented the positions of gene set members within the ranked gene list. Statistical metrics for the pathway are shown on the right. **(c)** Representative p21 immunostaining and quantification of p21+ cells per high-power field (HPF). Data were expressed as mean ± SEM; unpaired t-test. (**d)** Representative β-galactosidase staining and quantification of β-gal-positive area. Data were expressed as mean ± SEM; unpaired t-test. **(e)** Scatter plot showing Spearman correlation between normalized *Cdkn1a* and *Gpnmb* expression within IRI+PM2.5. Regression lines with 95% confidence interval (CI) were shown for visualization.

To determine whether inhaled PM2.5 particles could reach the kidney and directly contribute to the molecular changes observed in PT, we employed a combined scanning transmission electron microscopy–energy-dispersive X-ray spectroscopy (STEM–EDX) analysis in PT. STEM images visualize the ultrastructure of organelles, showing both mitochondria (Fig. 4f, HAADF image, red arrows), nuclear (Fig. 4f, HAADF image, blue arrow) and lysosomal compartments (Fig. 4f, HAADF image, black arrows). The EDX analysis enables the elemental identification and confirm the presence of PM2.5-associated elements at the ultrastructural level. Strikingly, elements detected in PT cells—including Al, Cl, and P—matched those identified in PM2.5 collected from Teflon filters near the exposure chamber. These elements were not diffusely distributed but accumulated within lysosomes (Fig. 4f; Supplementary Fig. S3f). This provides direct evidence of a full exposome trajectory-from air to kidney-demonstrating that components of inhaled PM2.5 can translocate to the kidney and accumulate within lysosomes of PT, where they may contribute to lysosomal stress and metabolic dysfunction.

### PM2.5-driven organelle stress promotes maladaptive repair through tubular senescence

The accumulation of PM2.5-derived elements within the PT lysosomes, together with mitochondrial injury and lipid dysregulation, suggests sustained toxic stress rather than a transient insult. Such metabolic and organelle stress are well-recognized triggers of cellular senescence and position PM2.5 as a driver of maladaptive repair, consistent with our prior findings^6^.

RNA-seq analysis identified *Cdkn1a* (p21) as the most upregulated gene in IRI+PM2.5 kidneys (Fig. 5a), highlighting cellular senescence as a major transcriptional response. Because p21 alone does not fully capture the senescence program, we performed GSEA using the SenMayo curated gene set, which confirmed significant enrichment of senescence pathways in IRI+PM2.5 kidneys (Fig. 5b). To determine whether PT display a senescent phenotype, we performed p21 IHC and senescence-associated β-galactosidase (SA-β-gal) staining. Both analyses showed a predominantly tubular localization and a marked increase in senescent cells in IRI+PM2.5 compared with IRI+FA kidneys (Fig. 5c,d).

Because senescent cells characteristically display lysosomal alterations^31^, we next examined whether PM2.5 induced lysosomal stress was linked to senescence induction in this setting. Supporting this idea, *Cdkn1a* expression positively correlated with *Gpnmb* (Fig. 5e), reinforcing the link between PM2.5-induced lysosomal stress and senescence.

### *In vitro* validation identifies nicotinamide as a metabolic rescue for PM2.5-challenged PTECs

To mechanistically validate our *in vivo* findings, we developed an *in vitro* model using immortalized proximal tubular epithelial cells (PTECs) subjected to Hypoxia/Re-oxygenation (H/R) injury^32^, followed by exposure to 20 µg/mL PM2.5 for three consecutive days (Fig. 6a). The PM2.5 used for these experiments was collected on Teflon filters placed adjacent to the exposure chamber. To determine which PM2.5 components drive PT damage, we selected filters with distinct profiles, including samples enriched in inorganic metals (metal-rich) and others relatively enriched in black carbon (BC-rich). Both components have been implicated in oxidative stress, mitochondrial injury, and tubular dysfunction in epidemiologic and experimental studies^16, 33–36^. Because BC cannot be reliably detected in kidney tissue by STEM–EDX, the use of component-enriched fractions provided a controlled setting to evaluate the contribution of individual PM2.5 constituents.

**Figure 6.**
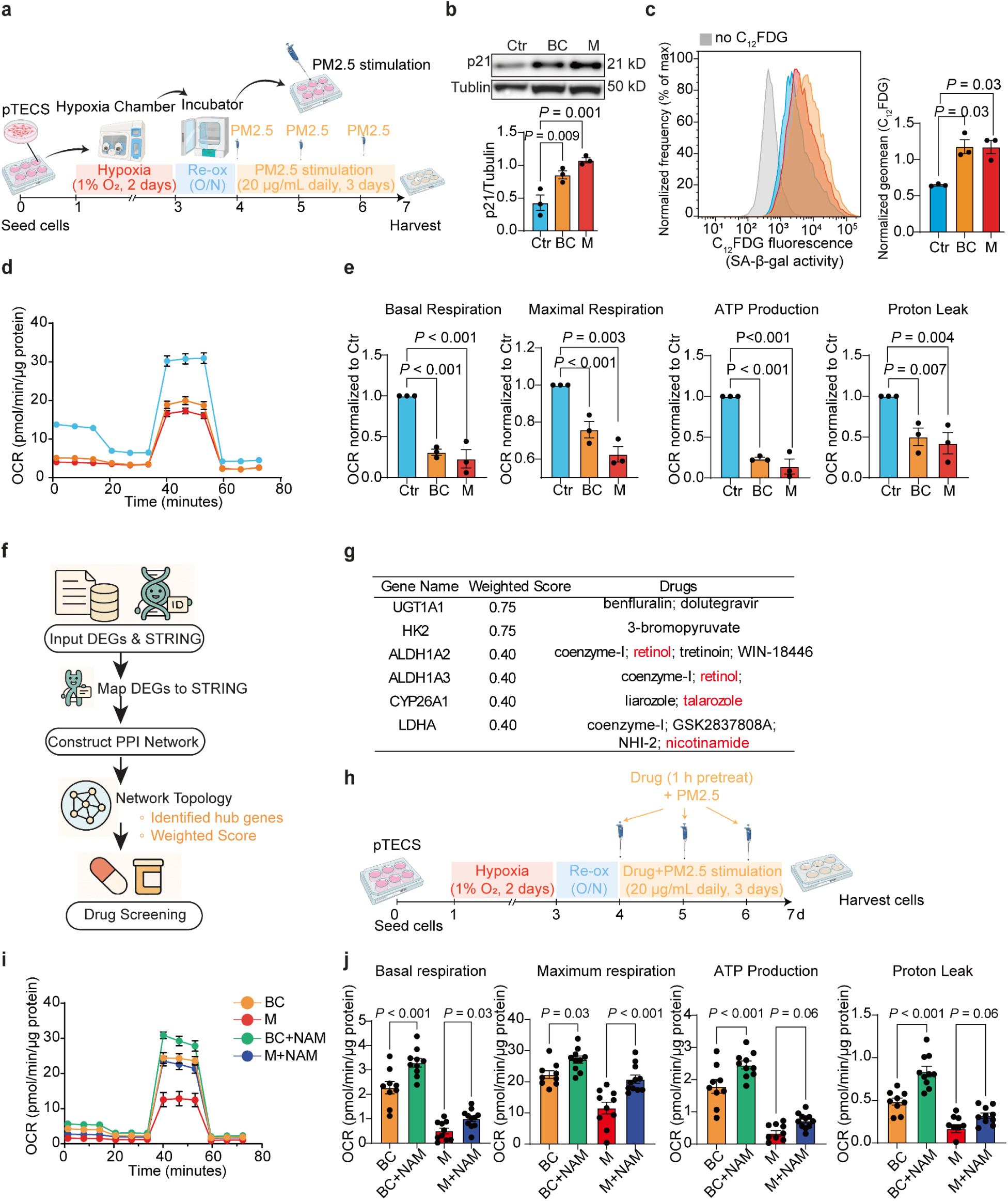
*In vitro* validation identifies nicotinamide as a metabolic rescue for PM2.5-challenged PTECs. **(a)** Experimental design for hypoxia/reoxygenation (H/R) and PM2.5 stimulation in proximal tubular epithelial cells (PTECs). Figure created with BioRender.com. **(b)** Western blot analysis and quantification of p21 expression normalized to tubulin. Experimental groups included Ctr (control), BC (PM2.5 enriched in black carbon), and M (PM2.5 enriched in metals). Data were expressed as mean ± SEM; one-way ANOVA. **(c)** Flow cytometry of senescence-associated β-galactosidase (SA-β-gal) activity using C₁₂FDG. Mean fluorescence intensity normalized to control. Data were expressed as mean ± SEM; one-way ANOVA. **(d–e)** Seahorse mitochondrial stress test. (d) Representative oxygen consumption rate (OCR) traces from Ctr, BC, and M groups. OCR values were normalized to protein content. (e) Quantification of basal respiration, maximal respiration, ATP production, and proton leak. Data were expressed as mean ± SEM; one-way ANOVA. OCR values were normalized to protein content and then to the control mean. **(f)** Workflow of drug screening for repurposing analysis. **(g)** Hub-gene network analysis and predicted interacting compounds; nutrient-like candidates (retinol, talarozole, nicotinamide(NAM)) highlighted in red. **(h)** Drug-intervention workflow to test the nutrient-like candidates. Figure created with BioRender.com. **(i-j)** Seahorse analysis of NAM rescue effects. (i) Representative OCR traces. OCR values were normalized to protein content. (j) Quantification of mitochondrial parameters. OCR values were normalized to protein content. Data were expressed as mean ± SEM; one-way ANOVA.

Both metal-rich or BC-rich–treated PTECs showed increased p21 protein expression following H/R injury (Fig. 6b), accompanied by elevated SA-β-gal activity(Fig. 6c), indicating induction of cellular senescence consistent with our *in vivo* observations. Similarly, combined H/R and exposure to metal-rich or BC-rich PM2.5 induced mitochondria dysfunction, with marked reductions in basal respiration, maximal respiratory capacity, ATP-linked respiration, and proton leak (Fig. 6d-e).

To further investigate the tubule–endothelial crosstalk observed *in vivo*, and specifically the influence of senescent PTECs, we employed a transwell co-culture system. PTECs were placed in the lower chamber, while endothelial cells were cultured in the upper insert (Supplementary Fig. S4a). Endothelial cells exposed to soluble mediators released by senescent PTECs (part of the senescence-associated secretory phenotype induced by metal-rich or BC-rich PM2.5 exposure) showed upregulated α-SMA protein expression (Supplementary Fig. S4b), suggesting that senescent PTECs can drive endothelial reprogramming consistent with EndMT observed *in vivo*.

Given these translational parallels between *in vivo* and *in vitro*, we next sought to identify potential molecular targets capable of mitigating PM2.5-induced mitochondrial and metabolic dysfunction. An integrated network analysis of DEGs was conducted using STRING to generate protein–protein interaction (PPI) networks. Network topology assessment yielded a set of hub genes that were prioritized as candidate targets for further investigation (Fig. 6f). We prioritized agents that act as nutrient-like metabolic supports rather than classical pharmacological modulators, as such compounds are easier to supplement and have a favorable safety profile. Based on prior evidence linking these compounds to kidney injury and repair^37, 38^, we focused on nicotinamide (NAM), retinol and talarozole (Fig. 6g). Cell viability assays confirmed that retinol and talarozole were non-toxic at the selected concentrations of 2 µM (Supplementary Fig. S4c), while NAM was administered at 1 mM as previously established in our group^39^. In this experimental setup, PTECs were subjected to H/R and, during the reoxygenation phase, pretreated for 1 hour with retinol, talarozole, or NAM before being exposed to metal-rich or BC-rich PM2.5 (Fig. 6h). Drug and PM2.5 were continuously present in the culture medium and treated daily for three consecutive days. While 2 µM retinol and talarozole failed to improve mitochondrial function following metal-rich or BC-rich PM2.5 exposure (Supplementary Fig. S4d), NAM preserved mitochondrial function based on respirometry assay (Fig. 6i-j), highlighting its protective effect.

## Discussion

Previously, we demonstrated that high levels of PM2.5 pre-exposure aggravated IRI-induced AKI^6^. Here, we show that even low-level ambient PM2.5 is sufficient to impair kidney recovery after injury. Although conventional kidney function remained largely preserved, integrated AI-based morphometrics, ultrastructural analyses, and multi-omics analysis revealed consistent molecular, metabolic, and microstructural abnormalities characteristic of maladaptive repair and premature kidney aging. These alterations included interstitial fibrosis, inflammation, microvascular and podocyte injury, PT damage characterized by mitochondrial dysfunction and metabolic rewiring, lipotoxicity and increased tubular senescence (Fig. 7). Reduced weight gain, a recognized predictor of adverse outcomes after AKI^40–43^, further reflected a sustained systemic stress state. We demonstrated that PM2.5-associated components were detected within PT lysosomes by EDX, indicating that inhaled particles can accumulate in the kidney after injury. Although the precise route of entry remains unclear, one plausible explanation is that particles may pass through the glomerular filtration barrier and reach the tubular compartment. Supporting this, we observed podocyte alterations, including a decrease in FSD, indicating an expansion of the foot process area and consistent with foot process effacement^18, 44^. Thus, podocyte injury could be a direct consequence of particle passage across the glomerular barrier. In parallel, PM2.5-induced microvascular rarefaction likely perpetuates cortical hypoxia, consistent with the dysregulation of hypoxia-responsive genes and compensatory VEGF overexpression. This phenotype echoes findings in developmental exposure studies^45^, where maternal PM2.5 exposure disrupts angiogenic signaling and reduces glomerular and peritubular endothelial, establishing a tissue environment that sustains hypoxia and increases susceptibility to further injury.

**Figure 7.**
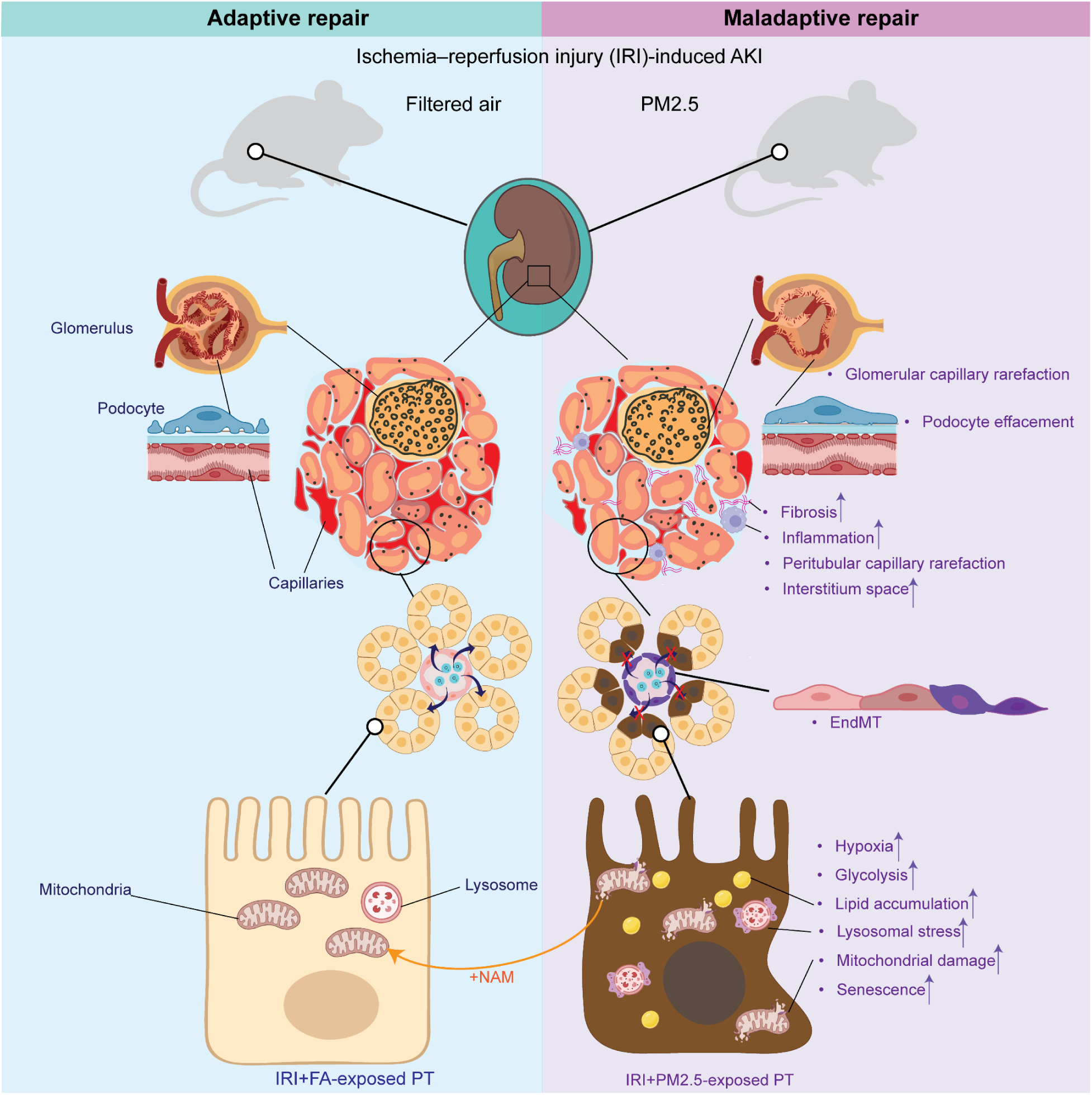
Summary of PM2.5-associated impairment in kidney recovery after ischemia–reperfusion injury (IRI). This illustration summarized the chronic structural and cellular alterations induced by PM2.5 exposure that impair kidney recovery after IRI. Key features included glomerular and peritubular capillary rarefaction, podocyte effacement, expanded interstitium space, interstitial fibrosis, inflammation, endothelial-to-mesenchymal transition (EndMT), increased tubular hypoxia, enhanced glycolytic activity, lipid accumulation, lysosomal stress, and elevated mitochondrial damage, culminating in the induction of tubular senescence. *In vitro*, administration of nicotinamide (NAM) restored mitochondrial function. Figure created with BioRender.com.

PTs have a high metabolic demand, relying on intact mitochondrial function, and the continuous endocytic activity makes these cells exceptionally vulnerable to injury from combined ischemic and particulate insults^7, 46^. Consistent with this intrinsic susceptibility, PM2.5 has been shown to induce EMT^47^ and apoptosis^7^ in human PTECs *in vitro*, and has been linked PT damage in humans^16^, aligning well with our observations. Indeed, we observed prominent tubular expression of senescence markers such as p21 and SA-β-gal. Importantly, this senescent phenotype was reproduced *in vitro*, demonstrating that PM2.5 exposure following hypoxia–reoxygenation is sufficient to induce tubular senescence independently of systemic factors. Furthermore, senescent pTECs promoted endothelial dedifferentiation in a transwell co-culture system, suggesting that tubular senescence may propagate microvascular dysfunction through paracrine signaling. This observation is consistent with that endothelial dedifferentiation is as a potential contributor to maladaptive repair and interstitial fibrosis following AKI to CKD progression^48–50^.

Since EDX analysis revealed PM2.5-elemental signatures of Al, Cl, and P localized within PT lysosomes, this suggests a endocytic uptake and trafficking of internalized PM2.5 into the degradative compartment^51^. Chronic exposure may exceed lysosomal clearance capacity, leading to lysosomal stress, a condition associated with aging-related lysosomal dysfunction^52^. Supporting this, lipidomic profiling revealed that PM2.5-exposed kidneys exhibited accumulation of lysosome-associated lipids including BMP, HexCer, and SM^53–55^. BMP, structural lipids enriched in endo-lysosomes, are well-established marker of impaired lysosomal degradation and accumulates across aging in the kidney and in CKD models^56–60^. Similarly, HexCer and SM are sphingolipids metabolically linked to ceramide^61^, and their accumulation has been associated with compromised lysosomal integrity and dysfunction^55, 62^. Notably, HexCer accumulates in aged kidneys and is further elevated in diabetic and metabolically stressed states^63, 64^. SM shows similar patterns in pathological kidney conditions^65–68^. In addition, sphingolipids are integral components of cellular membranes^69^. Changes in sphingolipid abundance are consistent with phospholipid membrane remodeling in senescent cells in response to metabolic stress and aging^70^. While lipidome remodeling is well documented in the acute phase after IRI^71^, lipid signatures distinguishing adaptive versus maladaptive repair after AKI remain poorly defined. Whether the sphingolipid signature observed here specifically reflects PM2.5-induced lysosomal dysfunction or a broader maladaptive repair signature this requires further investigation.

Consistent with lysosomal lipid alterations, we further observed a strong positive correlation between the expression of *Gpnmb*, a lysosomal transmembrane glycoprotein that is induced under lysosomal stress^72^, and the senescence marker *Cdkn1a,* supporting that PM2.5-induced lysosomal dysfunction contributes to tubular senescence. Notably, *Gpnmb* was previously identified as a pro-repair factor following ischemic kidney injury, where it promotes lysosome-dependent clearance of apoptotic debris through coordinated phagocytosis and autophagy, and is essential for effective tissue repair^73^. In addition to lysosomal alterations, pronounced mitochondrial damage characterized by calcium deposition, cristae disruption, matrix condensation, and swelling, was observed. Despite this damage, we did not detect evidence of FAO impairment but a shift toward glycolysis alongside upregulation of lipid catabolic pathways and prominent lipid accumulation, consistent with a compensatory and maladaptive metabolic response^27, 29^. As these findings are based on steady-state metabolomics, future flux-based studies are required to validate these metabolic shifts.

Finally, we explored potential therapeutic strategies to mitigate the PM2.5-induced maladaptive molecular response. To this end, we applied a transcriptomics-based drug-repurposing approach. Interestingly, the analysis highlighted several druggable target genes primarily linked to metabolic pathways. Among the prioritized candidates, we selected talarozole, retinol, and NAM. While retinol and talarozole have previously been implicated in tissue repair and regeneration^37^, neither restored mitochondrial function in our *in vitro* model. In contrast, NAM effectively rescued mitochondrial activity in both H/R- and PM2.5-exposed PTECs. As a precursor of NAD⁺, NAM is known to improve mitochondrial function, reduces oxidative stress and inflammation, and thereby limits maladaptive repair and fibrosis^38, 74–76^. NAM has also been reported to improve metabolic profiles in CKD patients, and increases NAD⁺ metabolites in humans with a favorable safety profile^77–79^. Importantly, NAM is evaluated in clinical settings of AKI, where a Phase 1 placebo-controlled study demonstrated that oral supplementation safely increased circulating NAD⁺ metabolites and was associated with reduced AKI, supporting its translational relevance^79^. However, given the pleiotropic effects of NAM on cellular metabolism, the precise mechanisms underlying its protective effects and the influence on LDHA in this context remain unclear.

In conclusion, this study provides experimental evidence that realistic, daily exposure to PM2.5 even at concentration within current limits, can impairs kidney recovery after IRI. By modeling post-injury exposure patterns that closely resemble real-life conditions, our findings highlight air pollution as an underrecognized but modifiable risk factor for kidney disease progression. These results have important public health implications, particularly for vulnerable populations such as individuals with prior kidney injury or reduced renal reserve and support the need for stronger environmental policies and preventive strategies to reduce chronic PM2.5 exposure and protect kidney health.

## Disclosure

All the authors declare no competing interests.

## Ethics approval statement

All procedures and numbers of animals were approved by the Institutional Animal Care and Use Committee (protocol no. 1589/2021) and carried out at the Faculty of Medicine of the University of São Paulo (FMUSP), Brazil.

## Acknowledgements

The authors would like to thank Tim and Nicole Endlich from Nipoka for the PEMP experiment and the Core Facility Metabolomics of the Amsterdam UMC (www.cfmetabolomics.nl) for the lipidomic and metabolomic analysis. Additionally, we would like to thank Dr. Sante Berlingerio for assistance in performing seahorse experiments and Jeannette Pankras for assistance in EM imagining.

## Author contributions

AT, LA, AR, JK, and SF designed the experimental strategy, interpreted the results and led the project. AT, AR and PS wrote the manuscript. PS, ACP, TRS, CFHW, LB, NC and HJB conducted the experiments. NvdW and IMS performed the EM and EDX analysis. GEJ and RHH contributed to the interpretation of lipidomics and metabolomics data. MdFA established and conceptualized the exposome chamber and provided the elemental and chemical characterization of PM2.5 samples. JJTHR provided feedback and advice on the pathological assessment. PB and MS contributed the pathomics analyses. All authors reviewed, revised, and approved the final version of the manuscript.

## Data statement

The data supporting the findings of this study are available within the manuscript and its Supplementary Information. Supplementary material is available online at <u>xxxxxx</u>. RNA-seq raw data and raw read counts generated during this study are publicly available in the Gene Expression Omnibus (GEO; accession number GSExxxxxx). Metabolomics and lipidomics datasets have been deposited in the MetaboLights repository (EMBL-EBI; accession number MTBLSxxxx). Additional data, methodological details, and analysis scripts are available from the corresponding author upon reasonable request.

