## Supplementary material for "Chronic exposure to low-concentration urban PM2.5 accelerates maladaptive repair after ischemic injury via mitochondrial dysfunction and lysosomal stress"

**Supplemental Material**

**Supplemental Figure S1.** GSEA enrichment analysis of Sham + PM2.5 vs. Sham + FA kidneys using the Hallmark gene set database.

**Supplementary Figure S2.** Chronic PM2.5 exposure exacerbated proximal tubular hypoxia, mitochondrial damage, and metabolic rewiring after AKI.

**Supplementary Figure S3.** Chronic PM2.5 after AKI drives proximal tubular lipid dysregulation and lysosomal element deposition.

**Supplementary Figure S4.** PM2.5 exposure after H/R disrupts endothelial–epithelial crosstalk and the effects of retinol and talarozole *in vitro*.

**Supplementary methods**

**Supplementary Table S1.** Kidney function parameters measured in metabolic cages.

**Supplementary Table S2.** Pathomics quantitative morphometric analysis for compartment-specific blocks and for global features.

**Supplementary Table S3.** Differential expression analysis by RNAseq.

### Supplementary Figure S1

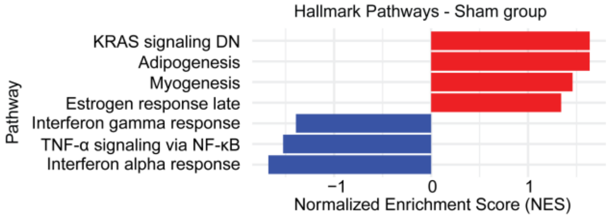


**Supplementary Figure S1.** Gene Set Enrichment Analysis (GSEA) of Hallmark pathway analysis showing significantly enriched pathways ($P < 0.05$) in Sham + PM2.5 vs. Sham + filtered air (FA) kidneys. Pathways were ranked by normalized enrichment score (NES).

### Supplementary Figure S2

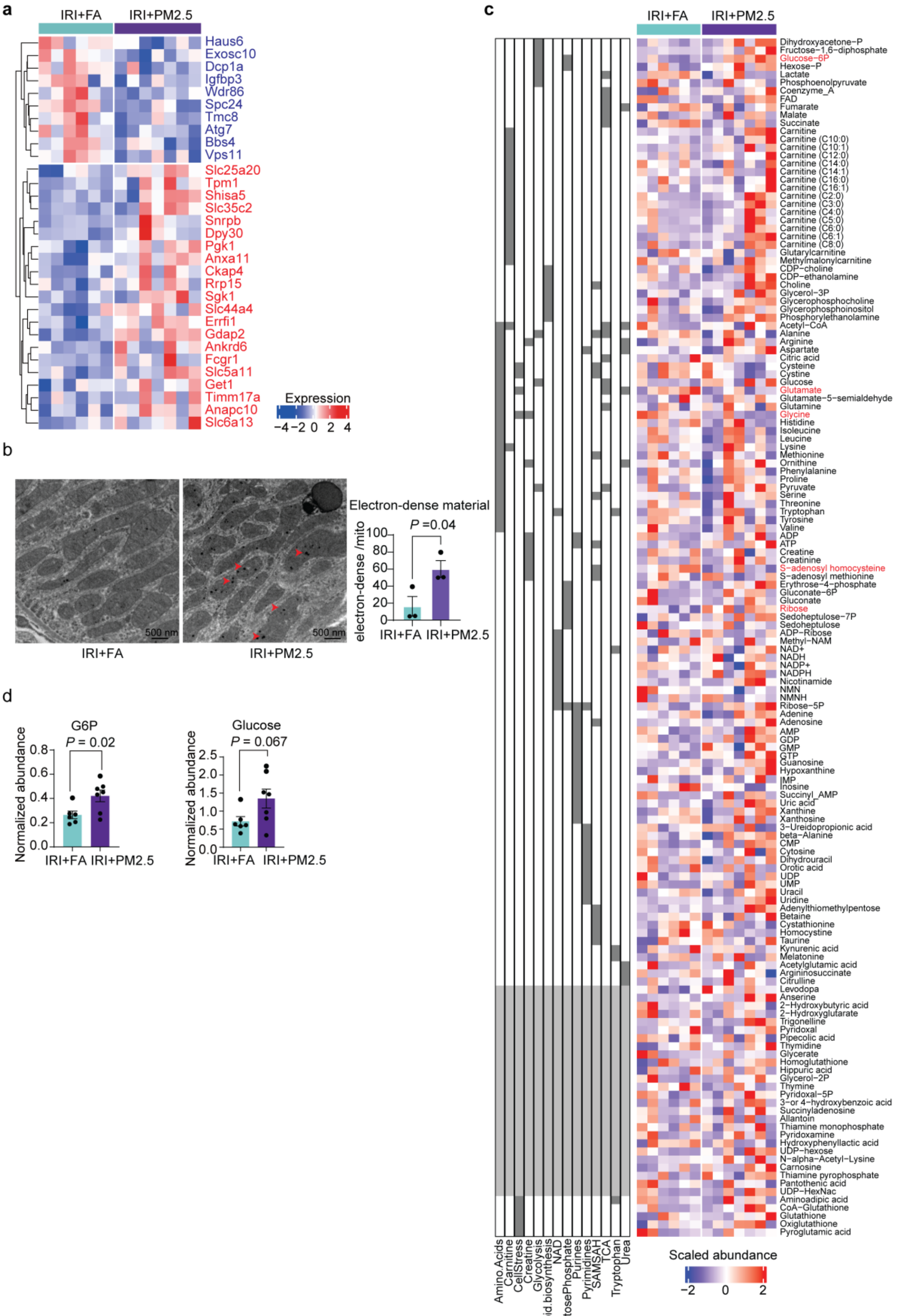


19

**Supplementary Figure S2. Chronic PM2.5 exposure exacerbated proximal tubular hypoxia, mitochondrial damage, and metabolic rewiring after acute kidney injury (AKI). (A)** Heatmap of 31 hypoxia-associated genes based on overlap with the iHypoxia database. 21 genes were upregulated (red labels) and ten downregulated (blue labels) in IRI+PM2.5 kidneys. **(B)** Electron-dense material normalized to mitochondrial number. Red arrowheads indicated electron-dense material within mitochondria. Data were presented as mean ± SEM; unpaired t-test. **(C)** Heatmap of scaled metabolite abundance (Z-score). Columns represented individual samples, and rows represent metabolites. The left annotation panel showed pathway assignments; grey boxes indicated pathway membership, whereas white or light grey boxes indicated no assignment. Metabolites with significant differences ($P < 0.05$) were labeled in red. **(D)** Normalized abundance of glucose-6-phosphate (G6P) and glucose from metabolomic analysis. Data were expressed as mean ± SEM; unpaired t-test.

### Supplementary Figure S3

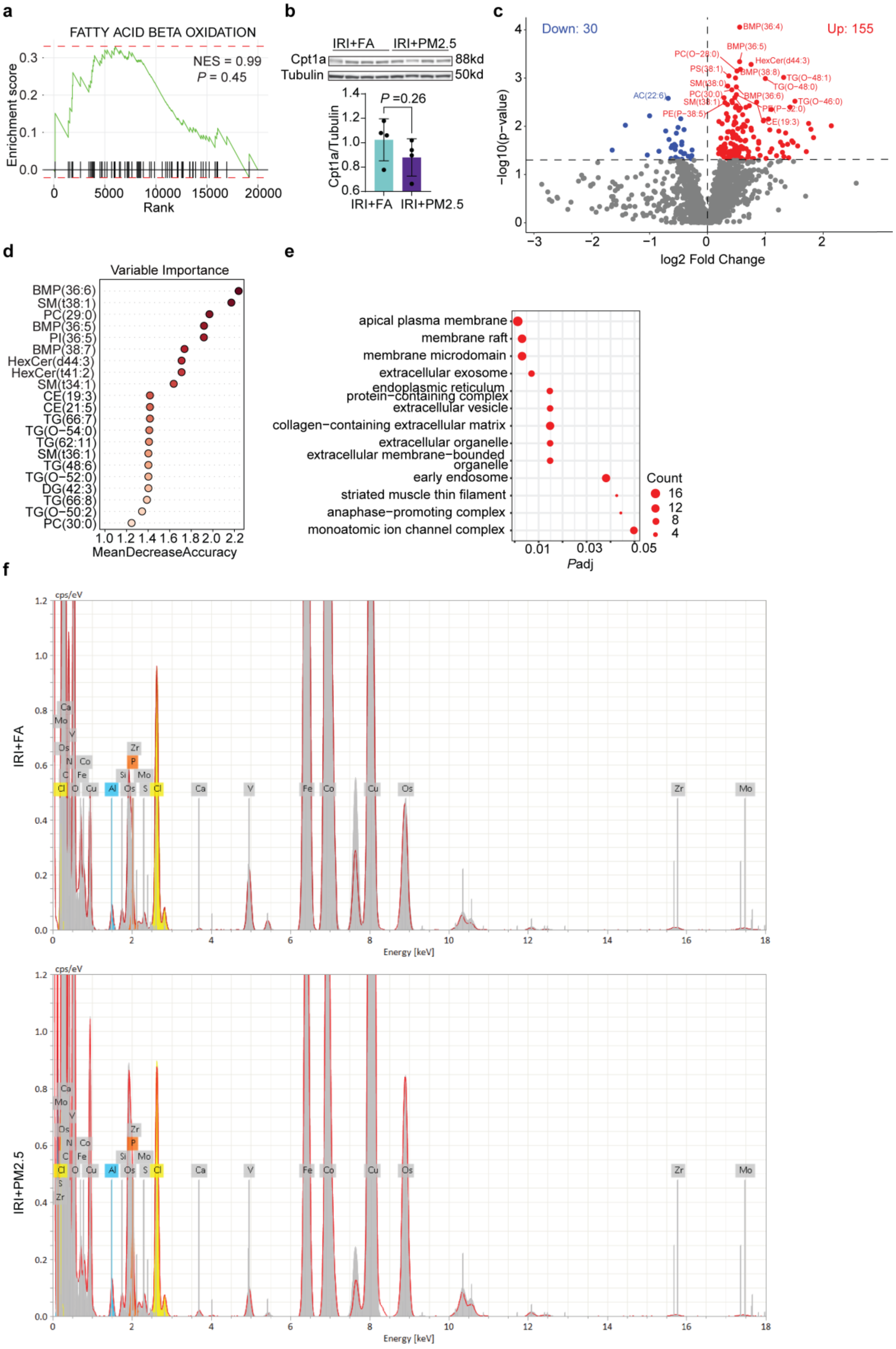

**Supplementary Figure S3. Chronic PM2.5 after acute kidney injury (AKI) drives proximal tubular(PT) lipid dysregulation and lysosomal element deposition.** **(A)** Gene Set Enrichment Analysis (GSEA) of fatty-acid β-oxidation from Gene Ontology–Biological Process (GO-BP). Green line indicated enrichment score; black vertical lines represented the positions of gene set members within the ranked gene list. Statistical metrics for the pathway were shown on the right. **(B)** Western blot and quantification of carnitine palmitoyltransferase 1A (Cpt1a). Data were expressed as mean ± SEM; unpaired t-test. **(C)** Volcano plot of differentially abundant lipid species. Significantly increased and decreased lipids ($P < 0.05$) were shown in red and blue, respectively. **(D)** Variable-importance ranking from random forest analysis, presented as mean decrease in accuracy; higher values indicated greater discriminatory contribution. **(E)** Gene Ontology - Cellular Component (GO-CC) enrichment of upregulated genes. Dot size indicated gene count. **(F)** Energy-dispersive X-ray (EDX) spectra of renal tissue after ischemia–reperfusion injury (IRI) with and without PM2.5 exposure. The y-axis showed counts per second per eV (cps/eV); the x-axis showed energy (keV). Element-specific peaks identified by the acquisition software were annotated. Representative spectra were acquired from electron-dense lysosomal inclusions in PT.

### Supplementary Figure S4

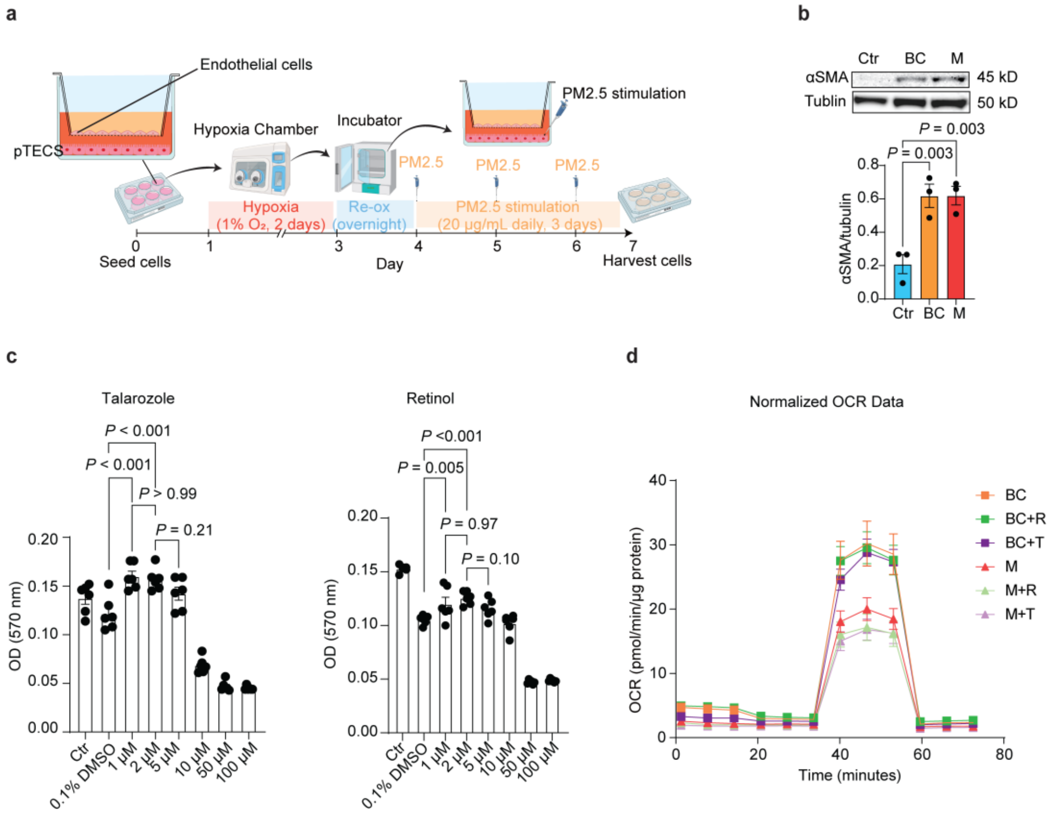


**Supplementary Figure S4. PM2.5 exposure after hypoxia–reoxygenation (H/R) disrupted endothelial–epithelial crosstalk and the effects of retinol and talarozole *in vitro*. (A)** Experimental schematic of the transwell co-culture model. Figure created with BioRender.com. **(B)** Representative western blot and quantification of α-smooth muscle actin (α-SMA) expression in endothelial cells under control (Ctr), baseline condition (BC), and PM2.5-treated (M) conditions. Tubulin was used as a loading control. Data were presented as mean ± SEM; one-way ANOVA. (**C)** Cytotoxicity of talarozole and retinol in proximal tubular epithelial cells (PTECs) was assessed using the MTT assay. 0.1% DMSO (corresponding to the 10 μM condition) was used as the vehicle control. Data were expressed as mean ± SEM; one-way ANOVA with Bonferroni correction. **(D)** Seahorse oxygen consumption rate (OCR) analysis in PM2.5 + retinol/talarozole groups. Data were expressed as mean ± SEM. The y-axis represented OCR normalized to protein content.

### Supplementary methods

#### Animals and study design

Male C57BL/6J mice (20–23 g), specific pathogen-free (SPF), were obtained from the animal facility of the School of Medicine of the University of São Paulo (FMUSP). They were housed under controlled environmental conditions (temperature, humidity, and a 12-hour light/dark cycle) with ad libitum access to food and water. All animal experiments were conducted in accordance with the ARRIVE (Animal Research: Reporting of In Vivo Experiments) guidelines and complied with the National Institutes of Health Guide for the Care and Use of Laboratory Animals. The study carried out at FMUSP, located at Av. Dr. Arnaldo, 455, Pacaembu, São Paulo, Brazil.  Mice were randomly assigned to four experimental groups: Sham + filtered air (FA) (n = 5), Sham + PM2.5 (n = 6), ischemia–reperfusion injury (IRI) + FA (n = 6), and IRI + PM2.5 (n = 7).

#### Surgical model of kidney IRI

Renal bilateral IRI was induced under ketamine (90 mg/kg) and xylazine (10 mg/kg, i.p.) anesthesia via a midline laparotomy. Kidney pedicles were clamped for 30 min with micro-clamps, followed by reperfusion for 6 months. Sham animals underwent identical procedures without vascular occlusion. Body temperature was maintained with controlled heating. Postoperative analgesia was provided with morphine sulfate (10 mg/kg, s.c.).

#### Whole-body exposure and tissue collection

Twenty-four hours after surgery, mice were housed in whole-body PM2.5 or FA exposure chambers from November 2021 to May 2022, located near FMUSP, which is located near downtown São Paulo, where air pollution is strongly influenced by emissions from both passenger and heavy-duty vehicles. PM2.5 or FA chambers were two independent but otherwise identical to environmental exposure conditions. In the PM2.5 chamber, ambient air was passed through an impactor to remove coarse particles (>2.5 µm), allowing continuous exposure to PM2.5, whereas in the FA chamber the same ambient air was filtered through a high-efficiency particulate air (HEPA) filter. An animation illustrating the whole-body exposure chamber setup is available at: https://www.pmkidney.com/educational.

Renal function and electrolyte homeostasis were assessed longitudinally by measuring serum and urine sodium and potassium levels, creatinine concentration, urinary osmolality, and 24-hour urinary sodium and potassium excreted loads. Plasma cystatin C levels were quantified as an additional marker of renal function. Detailed biochemical analyses were described in the Biochemical Analysis.

At the experimental endpoint, mice were anesthetized with ketamine (90 mg/kg) and xylazine (10 mg/kg, i.p.) and euthanized by exsanguination via cardiac puncture. Following euthanasia, kidneys were excised and rinsed in ice-cold saline. For histological analyses, tissues were fixed in 10% neutral-buffered formalin. Samples designated for ultrastructural analysis were fixed in EM grade fixative (0.1 M Phosphate buffer, 3% PFA, and 3% GA)  according to standard EM protocols. Tissue intended for cryosectioning was embedded in optimal cutting temperature (OCT) compound and stored at −80 °C until sectioning. All samples were stored under appropriate conditions until further processing.

**Biochemical Analysis**

Monthly, animals are placed in metabolic cages for 24 h urine and blood collection and body weight measurement. Urine and blood samples were aliquoted and centrifuged for 30 min at 4,000 × g. Serum and urine levels of sodium and potassium were quantified with an automated electrolyte analyzer (EasyLyte; Medica Corporation, Bedford, MA, USA). Creatinine concentration was determined with a commercial kit (Labtest Diagnóstica AS, Lagoa Santa, Brazil). Urinary osmolality was measured using a freezing-point depression osmometer (e.g., Advanced Instruments, Norwood, MA). Plasma samples were analyzed in duplicate according to the manufacturer's instructions. Briefly, a defined volume of plasma was loaded into the osmometer sample holder, and osmolality was determined based on the depression of the freezing point relative to pure water.

The osmometer was calibrated using manufacturer-provided standard solutions of known osmolality prior to sample analysis. Quality control samples were included to ensure measurement accuracy and instrument stability. Osmolality values were expressed as milliosmoles per kilogram of water (mOsm/L $H_2O$).

Urinary sodium and potassium excreted loads were calculated to estimate total electrolyte excretion during the urine collection period. The excreted load was calculated according to the following formula:

$$\text{Excreted Load}_{\text{Na or K}} = U_{\text{Na or K}} \times V_{/1000}$$

where $U_{\text{Na}}$ and $U_{\text{K}}$ represent urinary sodium and potassium concentrations, respectively, and V denotes the total urine volume collected over 24 hours in metabolic cages.

Plasma cystatin C levels were quantified using a commercially available sandwich ELISA kit specific for mouse cystatin C (DuoSet ELISA Development System, R&D Systems, Minneapolis, MN; Catalog #DY1238), according to the manufacturer's instructions. Briefly, 96-well microplates were coated overnight at room temperature with a capture antibody diluted in phosphate-buffered saline (PBS). Plates were washed and blocked with 1% bovine serum albumin (BSA) in PBS prior to sample incubation.

Plasma samples and standards were assayed in duplicate and diluted in reagent diluent (1% BSA in PBS) as appropriate. Following a 2-hour incubation at room temperature, plates were washed and incubated with a biotinylated detection antibody for 2 hours. After additional washing steps, streptavidin–horseradish peroxidase was added and incubated for 20 minutes, followed by color development using tetramethylbenzidine (TMB) substrate. The reaction was stopped with methanesulfonic acid, and absorbance was measured at 450 nm with wavelength correction at 540 or 570 nm using a microplate reader.

Standard curves were generated for each assay using a seven-point, two-fold serial dilution of recombinant mouse cystatin C. Optical density values were background-corrected by subtracting the zero-standard absorbance. Concentrations were calculated by fitting the data into a four-parameter logistic (4-PL) regression model. For diluted samples, values obtained from the standard curve were multiplied by the corresponding dilution factor. The assay exhibits high specificity, with no detectable cross-reactivity with related cathepsins or cystatin family members, as reported by the manufacturer.

**PM2.5 sampling and chemical characterization**

PM2.5 samples were collected over 24-hour periods from November 2021 to May 2022 using two Partisol 2025i Sequential Air Samplers (Thermo Fisher Scientific, USA) equipped with 47 mm Teflon filters (2 µm pore size; Pall Corporation, USA) and 47 mm quartz filters (QM-A; Whatman, UK). The samplers were positioned approximately five meters from the whole-body exposure chambers.

Teflon filters were conditioned for 48 hours under stable temperature and humidity prior to gravimetric weighing using a microbalance. After sampling, filters were reweighed and analyzed for trace elements (Na, Mg, Al, Si, P, S, Cl, K, Ca, V, Cr, Mn, Fe, Ni, Cu, Zn, As, Se, Br, Rb, Sb, and Pb) using an Epsilon 5 energy-dispersive X-ray fluorescence (ED-XRF) spectrometer (Malvern Panalytical, UK/The Netherlands).

Quartz filters were pre-heated at 600 °C for 6 hours in a muffle furnace to remove organic contaminants. Filters were weighed under controlled environmental conditions before and after sampling and subsequently stored at −20 °C until analysis. Organic carbon (OC) and elemental carbon (EC) concentrations were quantified using a Lab OC-EC Aerosol Analyzer (Sunset Laboratory, USA) based on the thermal-optical transmittance (TOT) method.

**Histology and Immunohistochemistry (IHC)**

Kidney tissues were fixed in 4% formalin at 4 °C for 24 h, dehydrated through a graded ethanol series, cleared in xylene, and embedded in paraffin. Sections (4 µm) were cut and deparaffinized for histological staining or immunohistochemistry (IHC).

For histology, Sirius Red staining for collagen was performed as previously described[S1] at the Department of Pathology, Leiden University Medical Center. Periodic acid–Schiff with diastase (PAS-D) staining was performed by incubating sections with 0.25% diastase (α-amylase) (Sigma-Aldrich, A3176-1MU) for 30 min at room temperature, followed by oxidation with 1% periodic acid for 10 min and staining with Schiff reagent (Merck; 109033) for 15 min.

For IHC, deparaffinized sections were rehydrated and treated with 3% hydrogen peroxide to block endogenous peroxidase activity. Antigen retrieval was performed in citrate buffer (pH 6.0) at 95 °C for 10 min. Sections were incubated overnight at 4 °C with primary antibodies against CD34 (CL8927ap; 1:1600), F4/80(Serotec MAC497 ga, 1:200) VEGF (Millipore, 07-1420; 1:200), PKM2 (Cell Signaling Technology, 4053; 1:5000), α-SMA (Dako, M851; 1:500), or p21 (Abcam, ab188224; 1:4000), all diluted in Normal Antibody Diluent (phosphate-buffered; CyTek Laboratories, ABB999). For α-SMA, rabbit anti-rat antibody (Southern Biotech, 6180-01) was applied prior to secondary detection. Secondary detection was performed using HRP-conjugated antibodies: goat anti-mouse (IL Immunologic, DPVM110HRP, The Netherlands) and goat anti-rabbit (IL Immunologic, DPVR110HRP, The Netherlands). Visualization was achieved using DAB, followed by hematoxylin counterstaining.

For α-SMA and CD34 double staining, CD34 (1:500) was detected using rabbit anti-rat (Southern Biotech, 6180-01), followed by goat anti-rabbit/polyAP and visualized with PermaBlue, whereas α-SMA (1:1000) was detected using goat anti-mouse/polyHRP and visualized with DAB. The resulting images were converted to an immunofluorescence format using ImageJ software (v1.54g, National Institutes of Health, USA). Negative controls were prepared by omitting the primary antibody.

Whole-slide images were digitized using a Philips IntelliSite Pathology Solution Ultra Fast Scanner (version 1.6) at a spatial resolution of 0.25 µm per pixel (40× equivalent magnification) and stored in pyramidal BigTIFF whole-slide image format.

**Pathomics analysis**

Whole-slide images of 4-µm PASD-stained kidney sections were digitized using a Philips IntelliSite Pathology Solution Ultra Fast Scanner (version 1.6) at a spatial resolution of 0.25 µm per pixel (40× equivalent magnification). Images were stored in pyramidal BigTIFF whole-slide image format. The digitized slides were subsequently analyzed using FLASH, a deep learning–based histomorphometry platform[S2].

Tissue compartments, including glomeruli, tubules, and arteries, were automatically segmented using instance segmentation. Quantitative morphometric features (e.g., size, diameter, and shape descriptors) were extracted at the level of individual segmented objects. To account for within-sample dependency among objects, linear mixed-effects models were fitted with experimental group as a fixed effect and sample as a random intercept. For area-based features, defined as the recognized feature area normalized to total tissue area, group differences were assessed using the Kruskal–Wallis test. Group-wise data are presented as mean ± SEM, and predefined pairwise comparisons are additionally reported.

For representative downstream analyses, measurements from all segmented objects within each tissue compartment were aggregated at the sample level, and the median value per feature was used. Partial least squares discriminant analysis (PLS-DA) was performed using sample-level median feature values to explore multivariate separation between experimental groups.

All quantitative results are reported in Supplementary Table S2.

**Transmission Electron Microscopy (TEM) and Energy Dispersive X ray (EDX) Microscopy**

Kidney tissue was collected and fixed in EM grade fixative (0.1 M Phosphate buffer, 3% PFA, and 3% GA) at room temperature (RT) and stored at 4 °C until processing. Fixed tissue was washed with distilled water and post-fixed in 1% osmium tetroxide (in $ddH_2O$) for 1 h. Samples were subsequently washed and dehydrated through a graded ethanol series (70%, 70%, 80%, 96%, 100%, 100%; 15 min each). Tissue was then infiltrated with a 1:1 mixture of Epon resin (Electron Microscopy Sciences) and propylene oxide and incubated overnight. The following day, the mixture was replaced with 100% Epon and incubated for 3 h at 37 °C, followed by polymerization at 65 °C for 2 days. Ultrathin sections were cut using a diamond knife (Diatome) on a Leica Ultracut UC7 ultramicrotome, and contrasted with uranyl acetate (UA Zero; Agar Scientific) and lead citrate (Electron Microscopy Sciences, cat. #22410). Samples were imaged with a JEOL JEM-1400 electron microscope operated at 80 kV, equipped with an Emsis Xarosa camera.

For elemental analysis by energy-dispersive X-ray spectroscopy (EDX), Epon-embedded tissue was sectioned into 100 nm slices using a diamond knife (Diatome) on a Leica Ultracut UC7 ultramicrotome. Sections were carbon-coated with a 15 nm layer using a Leica ACE600 sputter coater. EDX analysis was performed on a Talos L120C electron microscope operated at 120 kV in STEM mode, with the sample tilted 15° towards the Bruker XFlash 6T detector. Data were acquired using Esprit 2.2 software with an acquisition time of ~1 h per image. EDS spectra were analyzed, using standardless Cliff-Lorimer and series-fit, and based on the acquired information, illustrative STEM-EDS elemental maps were generated.

**TEM Mitochondrial Damage and Electron-Dense Material Scoring**

TEM was used to evaluate mitochondrial ultrastructural damage and the accumulation of electron-dense materials in PTECs. Only mitochondria within PTECs were included in the analysis. For each experimental group, three animals were analyzed. From each animal, five proximal tubules were randomly selected, and one representative micrograph per tubule was acquired at 4,000× magnification. Within each micrograph, all mitochondria were counted and classified as normal or damaged based on established ultrastructural criteria, including membrane integrity, matrix density, and cristae organization (Fig. 3D). The percentages of normal and damaged mitochondria were calculated for each tubule. Electron-dense materials were quantified and normalized to the total number of mitochondria per image. The mean value of the five analyzed tubules was used as the value for each animal in statistical analyses.

**Podocyte Exact Morphology Measurement Procedure (PEMP)**

PEMP was performed on 4 µm paraffin-embedded kidney sections mounted on adhesive slides to quantify podocyte morphology ( https://www.nipoka.com/ ). In brief, sections were immunofluorescent stained for the podocyte marker podocin (Ab), followed by whole-slide imaging and 3D structured illumination microscopy (3D-SIM). Podocyte regions were automatically analyzed with the PEMP algorithm, which results in quantitative readouts including filtration slit density (FSD) and filtration slit length (FSL), based on the analysis of 20 glomeruli per slide. Maximum intensity projections were generated, and additional image analysis was performed using ImageJ software (v1.54g, National Institutes of Health, USA).

**RNA-seq library preparation and sequencing**

Total RNA was extracted from fresh frozen mouse kidney tissue using TRIzol Reagent (Invitrogen, USA). RNA quality was assessed with an Agilent 2100 Bioanalyzer, and only high-quality RNA samples (RIN >7) were used for library preparation.

Sequencing libraries were constructed using the TruSeq Stranded mRNA LT Sample Prep Kit (Illumina, USA) according to the TruSeq Stranded mRNA Sample Preparation Guide (Part #15031047 Rev. E). Briefly, poly(A)+ mRNA was enriched, fragmented, and reverse-transcribed into cDNA. After end repair, 5’ and 3’ adapter ligation, and PCR amplification, the libraries were quality-checked and quantified.

Sequencing was performed on an Illumina platform (NovaSeq 6000) using paired-end 101 bp reads with Sequencing by Synthesis (SBS) technology. Raw base call (BCL) files generated by the Real Time Analysis (RTA) software were converted into FASTQ files using bcl2fastq without adapter trimming.

Raw sequencing reads in FASTQ format were first assessed with FastQC (v0.12.1) to evaluate base quality scores, GC content, sequence duplication levels, and overrepresented sequences. Adapter sequences and low-quality bases were trimmed using Trimmomatic (v0.39), and reads shorter than 75 bp after trimming were discarded. Clean reads were then mapped to the mouse reference genome GRCm39 (GENCODE release M30 annotation) using STAR (v2.6.1d) with default two-pass alignment mode to improve splice junction detection. Alignment quality metrics, including mapping rate, read distribution, and duplication rate, were assessed using Samtools (v1.18) and Qualimap (v2.2.2). Gene-level read counts were generated using featureCounts (v2.0.6) from the Subread package, using the GTF annotation file corresponding to GRCm39.

Lowly expressed genes were filtered by retaining only those with a raw read count ≥ 15 in at least one sample. Raw counts were normalized using DESeq2 (v1.40.2), and regularized log (rlog) transformation was applied to stabilize variance across the range of mean expression values[S3]. The transformed data were then used for principal component analysis (PCA) to assess the impact of IRI and PM2.5 exposure on gene expression. Subsequently, differential gene expression analysis was performed using DESeq2 (v1.40.2). Considering that our model was relatively mild and produced only moderate transcriptional alterations, a less stringent exploratory threshold ($P$< 0.05) was applied for identifying differentially expressed genes (DEGs) to increase sensitivity and capture biologically relevant signals that might otherwise be missed under stricter multiple-testing correction. For functional annotation, Gene Ontology (GO) enrichment analysis was performed using clusterProfiler (v4.14.4)[S4] on the identified DEGs, with significance defined as an adjusted $P$ value < 0.05 ($P$adj< 0.05). In addition, Gene Set Enrichment Analysis (GSEA) was conducted using a pre-ranked gene list ordered by $\log_2$ fold change ($\log_2$FC) values with 1,000 permutations; significance was defined as a nominal $P$ value < 0.05. Visualization was performed using ggplot2 (v3.5.2)[S5]. All analyses were conducted in R (v4.3.2).

**Drug screening and repurposing**

For drug screening and repurposing, DEGs were mapped to the STRING database (version 12.0, Mus musculus) to construct a protein–protein interaction network with a confidence score ≥0.6. Network topology was evaluated using degree, betweenness, and closeness centralities, which were standardized as z-scores and combined into a weighted score (0.5 × Degree + 0.3 × Betweenness + 0.2 × Closeness). Based on this ranking, the top hub genes were selected and subsequently compared with entries in the Repurposing Drugs Database (Broad Institute, 20200324) to identify potential compounds targeting these genes.

**LC-MS based lipidomics and metabolomics**

Lipidomics and metabolomics analysis were conducted only in IRI groups (IRI + FA, n = 6; IRI + PM2.5, n = 7). Frozen tissue sections (~20 mg) were collected on dry ice and stored at –80 °C until extraction. Lipidomics was performed as previously described[S6, S7] using an Ultimate 3000 binary HPLC system (Thermo Scientific) coupled to a Q Exactive Plus Orbitrap mass spectrometer (Thermo Scientific) operated in positive and negative ESI modes at the Metabolomics Facility of Amsterdam University Medical Center (UMC). Data were processed with a modified XCMS R pipeline, normalized to internal standards and sample mass, and analyzed by PLS-DA for multivariate analysis for univariate comparisons. Variable Importance in Projection (VIP) scores from the PLS-DA model estimated each lipid species' contribution to class separation, with VIP ≥ 1 considered important. The model was restricted to two latent variables to limit overfitting and complemented with univariate statistics. A random-forest classifier (seed = 415) with out-of-bag error estimation was used to assess feature importance by mean decrease accuracy. Lipid species with missing values were excluded from the model.

Metabolomics was performed as previously described[S7,S8]. Metabolites were separated by HPLC and analyzed by high-resolution mass spectrometry in both positive and negative electrospray ionization modes. Identification was based on exact mass and retention time, and abundances were normalized to internal standards (one per major metabolite class). The approach is semi-quantitative, and changes were interpreted in the context of consistent trends within metabolite classes. Annotated metabolites follow the Human Metabolome Database (HMDB; www.hmdb.ca). Data were analyzed using partial least squares discriminant analysis (PLS-DA) implemented in the mixOmics R package to identify latent variables maximizing covariance between metabolite profiles (X) and sample groups (Y), a method suited for high-dimensional and collinear datasets. Statistical differences were assessed on normalized data using unpaired t-tests, with $p < 0.05$ considered significant.

***In vitro* experiments**

Immortalized mouse proximal tubular epithelial cells (PTECs) were cultured in HK2 medium as previously described[S9], and immortalized mouse kidney endothelial cells were cultured in DMEM (Gibco) supplemented with 10% fetal bovine serum (FBS; Gibco), penicillin–streptomycin (P/S), and glutamine. Both cell lines were routinely maintained at 33 °C in the presence of interferon-γ (IFNγ) to sustain the temperature-sensitive SV40 large T antigen. One week before experiments, cells were shifted to 37 °C and cultured without IFNγ to inactivate the SV40 large T antigen and allow physiological growth. To induce hypoxia–reoxygenation (H/R) injury, PTECs were transferred to a hypoxia chamber (Whitley H35 Hypoxystation; 1% $O_2$, 5% $CO_2$, 37 °C) the day after seeding and maintained under hypoxic conditions for 2 days, followed by reoxygenation under normoxia conditions in a standard incubator overnight. PM2.5 particles were isolated from Teflon filters that collected airborne material every 24 h next to the exposome chambers at FMUSP, ensuring that the particles used for in vitro experiments were identical to those inhaled by the animals. These particles consisted predominantly of BC and metals (M). PM2.5 was dispersed in sterile water using an ultrasonic bath prior to application. Cells subjected to H/R were exposed to 20 µg/mL PM2.5 once daily for three consecutive days. Where applicable, PTECs and endothelial cells were co-cultured in a Transwell system (Corning®, 24 mm insert with 0.4 µm pore polyester membrane; Product No. 3450). Both cell types were subjected to hypoxic conditions, and PM2.5 exposure was applied to the PTECs during the co-culture.

Senescence of PTECs was assessed by SA-β-gal flow cytometry and Western blot detection of p21. For SA-β-gal analysis, cells were incubated with FVS780 viability stain (Invitrogen, Cat. No. 65-0856-14; final dilution 1:100,000), bafilomycin $A_1$ (Sigma-Aldrich, B1793; final concentration 100 nM), and $C_{12}$FDG (5-dodecanoylaminofluorescein di-β-D-galactopyranoside; Fisher Scientific, Product Code 11590276; final concentration 33 µM) and analyzed on a FACSCanto™ II cytometer. For western blot, PTECs were lysed in RIPA buffer (50 mM Tris pH7.5, 0.15 M NaCl, 2 mM EDTA, 1% deoxycholic acid, 1% NP-40, 4 mM sodium orthovanadate, 10 mM sodium fluoride), proteins were denatured under reducing conditions and separated on Bolt™ Bis-Tris Plus Mini Protein Gels (4–12%, 1.0 mm, WedgeWell™ format; Invitrogen, Cat. No. NW04127BOX), transferred onto PVDF membranes (Thermo Scientific™, Cat. No. 88518), and probed with anti p21 (Abcam, ab188224; 1:1000). Tubulin (Sigma-Aldrich, T0198 ;1: 20000) was used as loading control. Protein bands were visualized using an enhanced chemiluminescence (ECL) detection system (Thermo Scientific™ Pierce™ ECL Western Blotting Substrate, Cat. No. 32106), imaged with an Amersham ImageQuant 800, and quantified using ImageJ software (v1.54g, National Institutes of Health, USA).

Fatty acid oxidation in PTECs was assessed by Western blot detection of carnitine palmitoyltransferase 1A (Cpt1a; Abcam, ab128568; 1:1000). Endothelial-to-mesenchymal transition (EndMT) in kidney endothelial cells was evaluated by Western blot detection of α–smooth muscle actin (α-SMA; Dako, M851; 1:1000) under the same experimental conditions described above.

Mitochondrial function in PTECs was assessed using the Seahorse XF Cell Mito Stress Test Kit (Agilent Technologies, Lot: W09925) on a Seahorse XFe96 Analyzer (Agilent Technologies). Stimulated Cells were pre-incubated in a non-$CO_2$ incubator at 37 °C for 45 min Seahorse XF assay Medium (Agilent, Lot No.20623002) supplemented with 10 mM glucose (Agilent Technologies, Cat. No. 103577-100), 1 mM pyruvate (Agilent Technologies, Cat. No. 103578-100), and 2 mM glutamine (Gibco, Cat. No. 25030-024). Oxygen consumption rate (OCR) was measured under basal conditions and after sequential injections of oligomycin (1 µM), FCCP (1 µM), and rotenone/antimycin A (0.5 µM each). Data were analyzed using Wave software (Agilent Technologies) according to the manufacturer's instructions, with normalization to total protein content.

Cell viability was assessed using the MTT (Thiazolyl Blue Tetrazolium Bromide, Sigma-Aldrich, Cat. No. M2128) assay. Cells were seeded in 96 well plates, treated with the indicated stimuli and MTT (final concentration 1 mg/ml) was added during the last 45 minutes of the stimulation. The medium was carefully removed, and formazan crystals were dissolved in DMSO. Absorbance was measured at 570 nm using a microplate reader (Clariostar).

For compound testing, PTECs were subjected to H/R injury (2 days hypoxia followed by overnight reoxygenation) prior to drug application. Cells were pretreated with retinol (Sigma-Aldrich; #7632), talarozole (MedchemExpress; #115866), or nicotinamide (NAM, Sigma-Aldrich; #72340) one hour before PM2.5 exposure and maintained on the same regimen once daily for three consecutive days. Retinol and talarozole were applied at non-toxic concentrations of 1–2 µM, as determined by MTT assays, while NAM was supplemented at 1 mM based on previous published work [S10]. This dosing schedule ensured continuous compound availability throughout the exposure period. Control groups received vehicle (0.1% DMSO) at equivalent volumes.

**Statistical analysis**

All statistical analyses were performed using GraphPad Prism 10 (GraphPad Software, San Diego, CA). Data distribution was assessed with the Shapiro–Wilk test. For consistency across endpoints, data are presented as mean ± SEM. Statistical testing was performed using parametric or non-parametric methods as appropriate based on data distribution.

For normally distributed data, group differences were assessed using unpaired t-tests (for two-group experiments) or one-way ANOVA (for multi-group experiments), as specified in the figure legends. Non-normally distributed data were analyzed using non-parametric tests (Mann-Whitney test for two-group experiments or the Kruskal–Wallis test for multi-group experiments).

For animal experiments involving four groups, statistical testing was restricted to two a priori planned comparisons (Sham + FA vs. Sham + PM2.5 and IRI + FA vs. IRI + PM2.5) and *P* values are reported as nominal.

For cell experiments, group differences were assessed using one-way ANOVA. Predefined comparisons versus the control group were performed, and *P* values are reported as nominal unless otherwise specified in the figure legends.

A $P < 0.05$ was considered statistically significant.

**Supplementary Table S1. Kidney function parameters measured in metabolic cages.**

| Timepoint | Parameter | Units | Sham+FA mean ± SEM (n) | Sham+PM2.5 mean ± SEM (n) | IRI+FA mean ± SEM (n) | IRI+PM2.5 mean ± SEM (n) | *P* value (Sham: FA vs PM2.5) | *P* value (IRI: FA vs PM2.5) |
|---|---|---|---|---|---|---|---|---|
| **4 weeks** | Body Weight | grams | 22.2 ± 0.89 (5) | 22.9 ± 0.32 (6) | 17.9 ± 0.73 (6) | 17.8 ± 0.70 (7) | 0.48 | 0.91 |
| | Urinary Volume # | mL/Day | 0.76 ± 0.15 (5) | 1.1 ± 0.22 (6) | 1.5 ± 0.41 (6) | 2.1± 0.73 (7) | 0.37 | 0.92 |
| | Water Intake # | mL/Day | 4 ± 0.54 (5) | 4.3± 0.42 (6) | 5.8 ± 0.60 (6) | 7.1 ± 1.1 (7) | 0.69 | 0.55 |
| | Urinary Osmolality # | mOsm/L | 2480 ± 255.5 (5) | 2787 ± 244.1 (6) | 1441 ± 240.1 (5) | 1527 ± 364 (7) | 0.67 | 0.92 |
| | Sodium Excretion | mEq/Day | 0.12 ± 0.03 (5) | 0.13 ± 0.02 (6) | 0.11 ± 0.02 (6) | 0.10 ± 0.02 (7) | 0.64 | 0.96 |
| | Potassium excretion | mEq/Day | 0.31± 0.07 (5) | 0.41 ± 0.06 (6) | 0.28 ± 0.04 (6) | 0.27 ± 0.06 (7) | 0.26 | 0.85 |
| **9 weeks** | Body Weight # | grams | 24.1 ± 0.47 (5) | 24.1 ± 0.63 (6) | 23.7 ± 0.53 (6) | 22.6 ± 0.67 (7) | 0.84 | 0.29 |
| | Urinary Volume | mL/Day | 1.2 ± 0.15 (5) | 1.8 ± 0.13 (6) | 0.93 ± 0.13 (6) | 1.2 ± 0.21 (7) | 0.03* | 0.34 |
| | Water Intake # | mL/Day | 5.4 ± 0.6 (5) | 5 ± 0.37 (6) | 3.2 ± 0.48 (6) | 4 ± 0.31 (7) | 0.60 | 0.31 |
| | Urinary Osmolality | mOsm/L | 1900 ± 165.1 (5) | 1702 ± 241.9 (6) | 1919 ± 225.5 (6) | 2151 ± 238.6 (7) | 0.56 | 0.46 |
| | Sodium Excretion # | mEq/Day | 0.17 ± 0.02 (5) | 0.28 ± 0.03 (6) | 0.13 ± 0.02 (6) | 0.18 ± 0.02 (7) | 0.03* | 0.20 |
| | Potassium Excretion | mEq/Day | 0.28 ± 0.02 (5) | 0.61 ± 0.08 (6) | 0.28 ± 0.04 (6) | 0.37 ± 0.06 (7) | <0.001* | 0.24 |
| **14 weeks** | Body Weight | grams | 23.7 ± 0.45 (5) | 25.8 ± 0.54 (6) | 21.1 ± 0.66 (6) | 23.5 ± 0.63 (7) | 0.03* | 0.0086* |
| | Urinary Volume | mL/Day | 0.85 ± 0.22 (5) | 1.4 ± 0.15 (6) | 1.7 ± 0.32 (6) | 1.1 ± 0.15 (7) | 0.09 | 0.052 |
| | Water Intake | mL/Day | 5.8 ± 0.73 (5) | 3.5 ± 0.50 (6) | 4.8 ± 0.60 (6) | 4.6 ± 0.81 (7) | 0.036* | 0.78 |
| | Urinary Osmolality | mOsm/L | 2262 ± 47.3 (4) | 1782 ± 109.5 (6) | 1494 ± 161.7 (6) | 2118 ± 246.6 (6) | 0.085 | 0.016* |
| | Sodium Excretion | mEq/Day | 0.10 ± 0.01 (4) | 0.16 ± 0.02 (6) | 0.25 ± 0.04 (6) | 0.18 ± 0.03 (6) | 0.17 | 0.12 |
| | Potassium Excretion # | mEq/Day | 0.37 ± 0.06 (4) | 0.43 ± 0.05 (6) | 0.41 ± 0.06 (6) | 0.39 ± 0.06 (6) | 0.43 | 0.86 |
| **19 weeks** | Body Weight | grams | 27.0 ± 0.31 (5) | 27.4 ± 0.20 (6) | 26.7 ± 0.75 (6) | 25.6 ± 0.83 (7) | 0.68 | 0.21 |
| | Urinary Volume | mL/Day | 1.2 ± 0.15 (5) | 1.1 ± 0.31 (5) | 1.3 ± 0.20 (6) | 0.89 ± 0.16 (7) | 0.62 | 0.20 |
| | Water Intake | mL/Day | 6.6 ± 0.68 (5) | 6 ± 0.97 (6) | 4.5 ± 0.56 (6) | 5.7 ± 0.78 (7) | 0.61 | 0.26 |
| | Urinary Osmolality | mOsm/L | 1649 ± 143.2 (3) | 2470 ± 439.7 (5) | 1864 ± 107.6 (6) | 2095 ± 277.3 (7) | 0.11 | 0.54 |
| | Sodium Excretion | mEq/Day | 0.16 ± 0.04 (3) | 0.18 ± 0.06 (5) | 0.19 ± 0.04 (6) | 0.14 ± 0.02 (7) | 0.73 | 0.30 |
| | Potassium Excretion | mEq/Day | 0.41 ± 0.10 (3) | 0.48 ± 0.15 (5) | 0.44 ± 0.09 (6) | 0.29 ± 0.05 (7) | 0.69 | 0.26 |
| **23 weeks** | Body Weight | grams | 26.7 ± 0.65 (5) | 25.7 ± 0.58 (6) | 25.8 ± 0.62 (6) | 22.6 ± 0.70 (7) | 0.31 | 0.0015* |
| | Urinary Volume | mL/Day | 1.3 ± 0.17 (4) | 2.0 ± 0.25 (5) | 1.7 ± 0.27 (6) | 2.2 ± 0.29 (6) | 0.09 | 0.13 |
| | Water Intake | mL/Day | 5.3 ± 0.63 (4) | 7 ± 0.95 (5) | 6.8 ± 0.70 (6) | 8 ± 0.89 (6) | 0.19 | 0.30 |
| | Urinary Osmolality | mOsm/L | 2100 ± 247.4 (4) | 2102 ± 272.4 (6) | 1243 ± 268.5 (5) | 890.4 ± 142.8 (7) | 0.007 | 0.28 |
| | Sodium Excretion | mEq/Day | 0.22 ± 0.04 (4) | 0.30 ± 0.01 (5) | 0.17 ± 0.03 (5) | 0.18 ± 0.03 (6) | 0.07 | 0.78 |
| | Potassium Excretion | mEq/Day | 0.48 ± 0.096 (4) | 0.7 ± 0.04 (5) | 0.32 ± 0.07 (5) | 0.3 ± 0.06 (6) | 0.78 | 0.13 |

*Data are presented as mean ± SEM (n = number of animals per group). Due to missing values, n varied slightly across parameters.*
*# Statistical analyses of parameters were performed using non-parametric tests.*
** P value < 0.05 was considered statistically significant.*
*Abbreviations: IRI, ischemia–reperfusion injury; FA, filtered air*

### Supplementary methods reference


S1. van Aanhold CCL, Koudijs A, Dijkstra KL*, et al.* The VEGF Inhibitor Soluble Fms-like Tyrosine Kinase 1 Does Not Promote AKI-to-CKD Transition. *Int J Mol Sci* 2022; **23**.

S2. Holscher DL, Bouteldja N, Joodaki M*, et al.* Next-Generation Morphometry for pathomics-data mining in histopathology. *Nat Commun* 2023; **14:** 470.

S3. Love MI, Huber W, Anders S. Moderated estimation of fold change and dispersion for RNA-seq data with DESeq2. *Genome biology* 2014; **15:** 550.

S4. Wu T, Hu E, Xu S*, et al.* clusterProfiler 4.0: A universal enrichment tool for interpreting omics data. *The innovation* 2021; **2**.

S5. Wickham H. Data analysis. *ggplot2: elegant graphics for data analysis*. Springer, 2016, pp 189–201.

S6. Herzog K, Pras-Raves ML, Vervaart MA*, et al.* Lipidomic analysis of fibroblasts from Zellweger spectrum disorder patients identifies disease-specific phospholipid ratios. *J Lipid Res* 2016; **57:** 1447–1454.

S7. Molenaars M, Schomakers BV, Elfrink HL*, et al.* Metabolomics and lipidomics in Caenorhabditis elegans using a single-sample preparation. *Dis Model Mech* 2021; **14**.

S8. Schomakers BV, Hermans J, Jaspers YRJ*, et al.* Polar metabolomics in human muscle biopsies using a liquid-liquid extraction and full-scan LC-MS. *STAR Protoc* 2022; **3:** 101302.

S9. Sanches TR, Parra AC, Sun P*, et al.* Air pollution aggravates renal ischaemia-reperfusion-induced acute kidney injury. *J Pathol* 2024; **263:** 496–507.

S10. Chanvillard L, Lantermans H, Wall C*, et al.* NNMT promotes tubular senescence and fibrosis in chronic kidney disease. *bioRxiv* 2025**:** 2025.2001.2006.631437.
